# Extracellular Vimentin is a Damage-Associated Molecular Pattern Protein Serving as an Agonist of TLR4 in Human Neutrophils

**DOI:** 10.1101/2024.05.02.592157

**Authors:** Łukasz Suprewicz, Krzysztof Fiedoruk, Karol Skłodowski, Magdalena Zakrzewska, Alicja Walewska, Piotr Deptuła, Agata Lesiak, Sławomir Okła, Peter A. Galie, Alison E. Patteson, Paul A. Janmey, Robert Bucki

## Abstract

**Background:** Vimentin is a type III intermediate filament protein, that plays an important role in cytoskeletal mechanics. It is now known that vimentin also plays important roles outside the cell. Recent studies show the controlled release of vimentin into the extracellular environment, where it functions as a signaling molecule. Such observations are expanding our current knowledge of vimentin as a structural cellular component towards additional roles as an active participant in cell signaling.

**Methods:** Our study investigates the immunological roles of extracellular vimentin (eVim) and its citrullinated form (CitVim) as a damage-associated molecular pattern (DAMP) engaging the Toll-like receptor 4 (TLR4) of human neutrophils. We used *in vitro* assays to study neutrophil migration through endothelial cell monolayers and activation markers such as NADPH oxidase subunit 2 (NOX2/gp91phox). The comparison of eVim with CitVim and its effect on human neutrophils was extended to the induction of extracellular traps (NETs) and phagocytosis of pathogens.

**Results:** Both eVim and CitVim interact with and trigger TLR4, leading to increased neutrophil migration and adhesion. CitVim stimulated the enhanced migratory ability of neutrophils, activation of NF-κB, and induction of NET formation mainly mediated through reactive oxygen species (ROS)-dependent and TLR4-dependent pathways. In contrast, neutrophils exposed to non-citrullinated vimentin exhibited higher efficiency in favoring pathogen phagocytosis, such as *Escherichia coli* and *Candida albicans*, compared to CitVim.

**Conclusions:** Our study identifies new functions of eVim in its native and modified forms as an extracellular matrix DAMP and highlights its importance in the modulation of immune system functions. The differential effects of eVim and CitVim on neutrophil functions highlight their potential as new molecular targets for therapeutic strategies aimed at differential regulation of neutrophil activity in different pathological conditions. This, in turn, opens new windows of therapeutic intervention in inflammatory and immunological diseases characterized by immune system dysfunction, in which eVim and CitVim play a key role.

## Introduction

Neutrophils are the most abundant leukocytes in human circulation and play a vital role in the inflammatory process (1). Neutrophils migrate to the site of inflammation and release antimicrobial agents, pro-inflammatory cytokines, and a wide range of other factors (2). Neutrophil activation can be triggered by various external stimuli, including proteins classified as damage-associated molecular patterns (DAMPs). DAMPs are endogenous molecules that accumulate during cellular injury or stress and act as ‘alarmins’ or ‘danger’ signals in response to infectious and non-infectious pathologies (3, 4). DAMPs stimulate neutrophils by binding pattern recognition receptors (PRRs), including Toll-like receptors (TLRs). These receptors, often activated by aberrantly expressed or bioactive fragments of extracellular matrix components (ECM), serve as critical mediators in the DAMP-induced activation of neutrophils (5). The role of DAMPs in neutrophil activation represents a delicate balance. While they are critical in defending against pathogens, their excessive or dysregulated activation can lead to detrimental consequences (6–8).

Vimentin (Vim), an intermediate filament protein, is predominantly expressed in mesenchymal cells and forms part of the cytoskeleton, maintaining cellular integrity and facilitating migration (9, 10). New work is highlighting however that these intracellular vimentin filaments can be disassembled into subunits that are released into the extracellular environment, coined as extracellular vimentin (eVim), which can then attach to the cell surface or the ECM (11–13). Increased expression of eVim and its post-translationally modified (PTM) form, citrullinated vimentin (CitVim), has been observed in various acute inflammatory conditions such as sepsis and chronic ones like pulmonary fibrosis or rheumatoid arthritis (RA) (14–16). Citrullination is a modification mediated by peptidyl arginine deiminase (PAD) enzymes (17) that convert the amino acid arginine into citrulline, leading to protein structure and function changes (18). Neutrophils, for example, activate PADs to disassemble their vimentin cytoskeleton, as they also citrullinate histones to release their DNA to form neutrophil extracellular traps (NETs) (19). Citrullination occurs in the joints of individuals with RA and results in the production of autoantibodies against citrullinated proteins, including CitVim, which can trigger an autoimmune response, leading to chronic inflammation (20). However, the specific roles of eVim and CitVim in neutrophil-mediated inflammation and their potential as therapeutic targets remain largely unexplored.

Here, we present evidence that eVim activates neutrophils, and citrullination enhances the stimulatory effect of eVim. We conducted a series of biochemical assays, including atomic force microscopy and inhibition studies, to demonstrate the interaction between eVim and CitVim with TLR4, providing insights into the molecular mechanisms involved. Through a combination of cellular assays, microscopy, 3D microfluidic models, and gene expression profiling, we assessed the influence of eVim and CitVim on neutrophil activities, including adhesion, migration, phagocytosis, NET formation, and cytokine secretion. The presence of CitVim significantly enhances specific neutrophil activities over non-citrullinated vimentin, but eVim is superior in boosting phagocytic efficacy. These findings expand our understanding of the role of eVim and PTMs in neutrophil biology and suggest a promising therapeutic approach for the treatment of inflammatory diseases.

## Results

### Extracellular vimentin on endothelial cells and enhanced adhesion and activation of human neutrophils

We assessed eVim presence at the surface of human umbilical vein endothelial cells (HUVECs) (**Fig. 1**). Our immunofluorescence study revealed that in HUVECs, eVim, unlike its intracellular filamentous form, appeared as surface dots (**Fig. 1a**). Such a pattern is consistent with previous reports (21, 22). To rule out antibody-induced membrane permeability changes, we performed control experiments with the HUVECs using an anti-β-actin antibody, confirming no intracellular actin stress fiber during vital staining (**Fig. S1a**). As shown in **Fig. S1b**, HUVECs lack detectable CitVim on their surface, but exogenous CitVim can bind the cell surface when added to the cell culture medium (**Fig. 1b**). Furthermore, CitVim from neutrophil-derived NETs also bound to the endothelial cell surface (**Fig. 1b**). We tested whether eVim or CitVim impacts neutrophil adhesion to cells using a confluent monolayer of vimentin −/− mouse embryonic fibroblasts (mEFs) preincubated with eVim or CitVim (**Fig. 1c**). Analysis of cell surface attachment of eVim and CitVim to vim −/− mEFs, revealed that the eVim and CitVim adhered to the surface of the vim −/− mEFs, displaying a punctate pattern (**Fig. S2a, b**). We observed a dose-dependent increase in adherent neutrophils after the addition of eVim (**Fig. 1d**), with an approximately 6-fold increase in adhesion at an eVim concentration of 20 μg/mL. Addition of CitVim had an even greater effect on neutrophil adhesion, with an approximately 9-fold increase at a CitVim concentration of 20 μg/mL.

**Figure 1.**
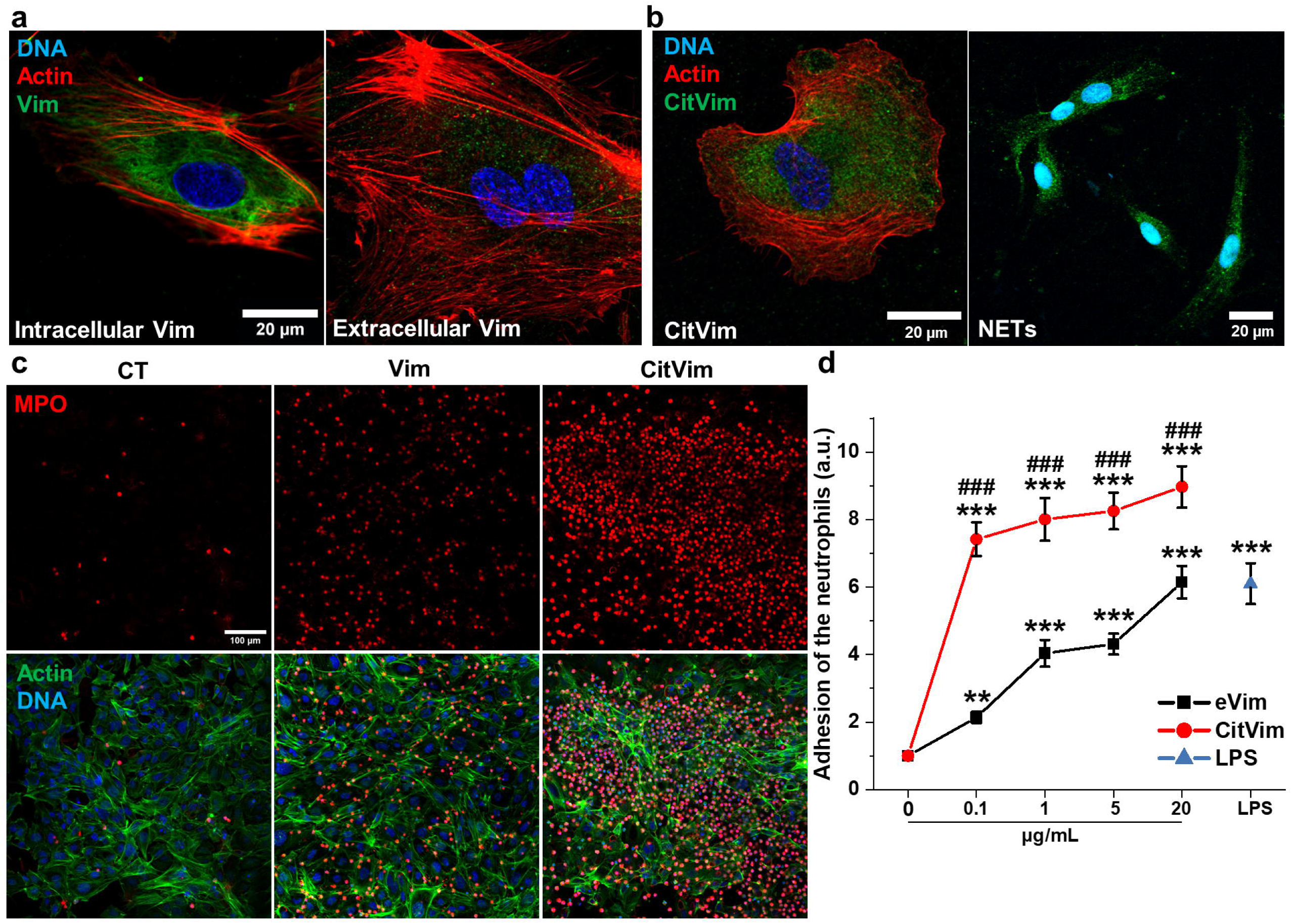
Extracellular vimentin triggers the activation and adhesion of human neutrophils. **a - b** Images of intracellular and extracellular vimentin (a) in human umbilical vein endothelial cells (HUVECs). Staining of citrullinated vimentin on the surface of HUVECs after the addition of citrullinated vimentin or supernatant from stimulated neutrophils containing NETs (b) for 1 h. Vimentin or citrullinated vimentin (green), actin (red), and nuclei (blue). Scale bar, ~20 µm. **c** Images of human neutrophils that adhered after 2 h of incubation to the monolayer of mouse embryonic fibroblasts lacking vimentin expression (vim−/− mEFs) preincubated or not with vimentin or citrullinated vimentin for 1 h. Neutrophils stimulated with LPS were used as a control. MPO – myeloperoxidase (red), actin (green), nuclei (blue). Scale bar, ~100 µm. **d** Quantification of calcein AM-stained neutrophils that adhere after 2 h to vim−/− mEFs monolayer preincubated with vimentin or citrullinated vimentin for 2 h (n = 3). LPS at 1 µg/mL was used as a control. Data are presented as the mean ± standard deviation of the mean. *, P ≤ 0.05; **, P < 0.01; ***, ###, P < 0.001. Hash (#) denotes statistical significance between corresponding conditions in citrullinated vimentin compared to vimentin (**d**). Significance was determined by one-way ANOVA with Tukey’s test.

To eliminate the potential influence of other proteins in our experimental settings, we fabricated 30 kPa polyacrylamide hydrogels functionalized with collagen (Col), eVim, or CitVim (**Fig. S3**). This method effectively isolated the impact of eVim and CitVim on neutrophil adhesion by eliminating other cell surface proteins, clarifying their specific roles (23). Both forms of Vim supported neutrophil adhesion, but more neutrophils adhered to the CitVim-coated hydrogels compared to eVim-coated hydrogels (**Fig. 2a**). The number of cells on the Col-coated surface was similar to that on the Vim-coated surface (**Fig. 2b**), but the average cell area on Vim-coated hydrogels was greater (**Fig. 2c**). Neutrophils on CitVim-coated hydrogels showed the largest cell area. Additionally, we observed neutrophils tend to form aggregates on both eVim and CitVim substrates with the highest number of cell clusters on CitVim-coated hydrogels (**Fig. 2d**). Moreover, we observed a significant increase in the NADPH oxidase subunit 2, NOX2/gp91phox (**Fig. 2e**), an indicator of the production of reactive oxygen species (ROS) and neutrophil activation, in neutrophils on both eVim and CitVim-coated hydrogels (24). These data suggest that eVim and CitVim, as a coating on hydrogels, enhance neutrophil activation, indicating a potential role of surface vimentin in the inflammatory response.

**Figure 2.**
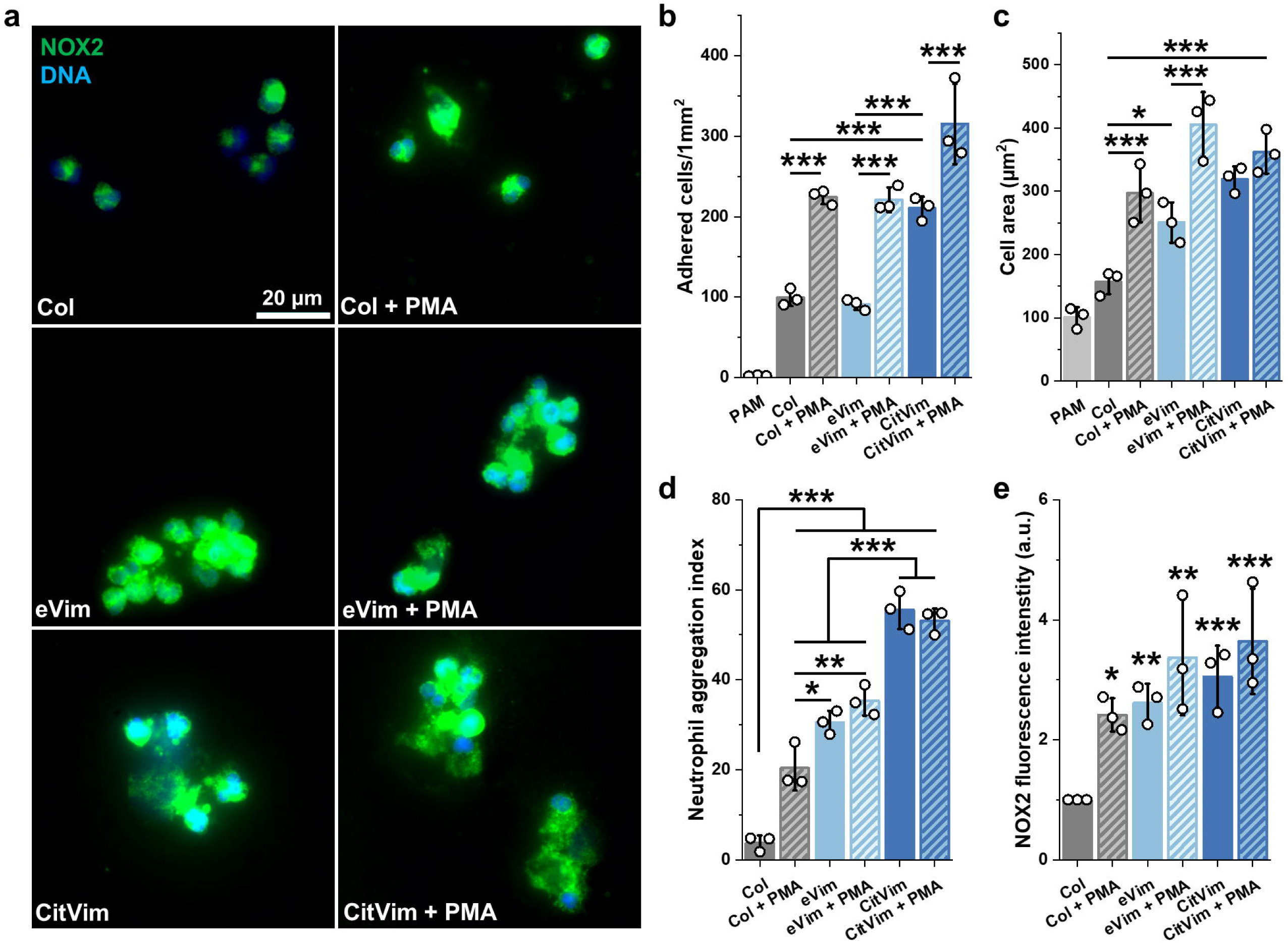
Extracellular vimentin activates human neutrophils when present on the surface of a polyacrylamide hydrogel. **a** Images of human neutrophils adhered on 30 kPa polyacrylamide hydrogels coated with collagen (Col), vimentin (eVim), and citrullinated vimentin (CitVim) at 0.1 mg/mL. Neutrophils were added to hydrogels for 2 h, then unbounded cells were washed off, captured for bright field imaging, and fixed for fluorescence staining. Phorbol 12-myristate 13-acetate (PMA) at 100 nM was used as a positive control. NOX2 (green), nuclei (blue). Scale bar, ~ 20 µm. **b - c** Number per 1 mm^2^ (b) and mean cell area of human neutrophils that adhere after 2 h of incubation to the 30 kPa hydrogels coated with collagen, vimentin, or citrullinated vimentin (n = 3). **d** Aggregation index of human neutrophils that adhere after 2 h of incubation to the 30 kPa hydrogels coated with collagen, vimentin, or citrullinated vimentin (n = 3). **e** Mean NOX2 fluorescence intensity in neutrophils seeded on Col, eVim, and CitVim coated hydrogels (n = 3). Results were normalized and presented relative to those of the collagen condition (Col), set as 1. Data are presented as the mean ± standard deviation of the mean. *, P ≤ 0.05; **, P < 0.01; ***, P < 0.001. Significance was determined by one-way ANOVA with Tukey’s test.

### Extracellular vimentin promotes transendothelial migration of human neutrophils

Since transendothelial migration is a crucial step in neutrophil extravasation at sites of infection or inflammation (25), we investigated the effects of vimentin on neutrophil migration in both transwell assays (**Fig. 3a - c**) and a more physiologically relevant three-dimensional (3D) microfluidic endothelial model (**Fig. 3d - h**). In the Transwell assay, more neutrophils adhered to eVim- or CitVim-coated membranes compared to membranes coated with collagen (**Fig. 3b**). This increased adhesion correlated with the number of cells that migrated through the pores of the membrane to the lower chamber (**Fig. 3c**). In the microfluidic model, a cylindrical void was created within a collagen hydrogel and seeded with HUVECs to form a confluent monolayer, simulating the blood vessel lining (**Figure 3d**). Subsequently, we introduced neutrophils stimulated with eVim, CitVim, or lipopolysaccharide (LPS) into the vessel, as well as unstimulated neutrophils as a control (**Fig. 3e**). LPS treatment of neutrophils was used for comparison with the effect of a potent neutrophil activator (26). eVim augments neutrophil adhesion within the blood vessels, and CitVim increases this effect (**Fig. 3f**). The number of neutrophils escaping the vessel lumen and the distance they traveled within the collagen hydrogel increased in the presence of eVim and was greater upon CitVim addition (**Fig. 3g, h**) compared to the control and LPS conditions. In comparison, non-stimulated neutrophils (CT) adhered poorly to the endothelial wall, and their extravasation rate was negligible.

**Figure 3.**
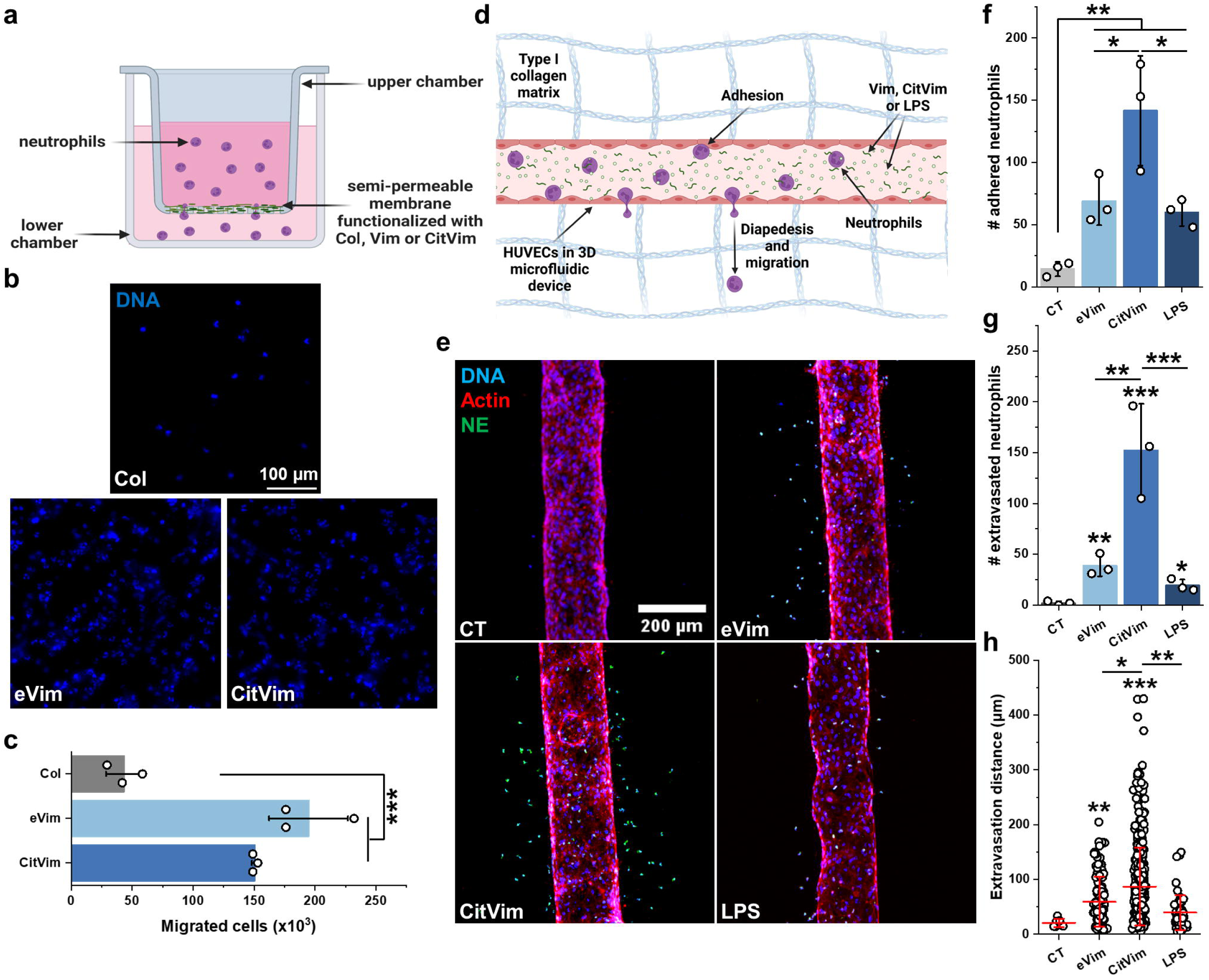
Extracellular vimentin promotes transendothelial migration of human neutrophils. **a** Schematic of the 2D human neutrophils migration model using Transwell inserts with 5 µm pore size. Each transwell insert was functionalized with collagen, vimentin, or citrullinated vimentin at 0.1 mg/mL. Unbounded proteins were washed, and human neutrophils were introduced to the upper chamber. After 2h, the upper chamber was washed and fixed for fluorescence imaging of neutrophils stuck in the pores or semi-permeable membrane. **b** Images present nuclei (blue) of neutrophils on the bottom side of the insert membrane. Scale bar, ~ 100 µm. Medium from the lower chamber was collected, and the number of migrated cells was counted (**c**) (n = 3). **d** Schematic of the 3D microfluidic vascular model of transendothelial migration of human neutrophils. HUVECs were grown in cylindrical channels (180 µm in diameter) in a type I collagen matrix. When endothelial cells reached full coverage, 2×10^4^ human neutrophils untreated or treated with 1 of vimentin, citrullinated vimentin, or LPS (all at 1 µg/mL) were introduced to the lumen for 2 h. After 2 h, channels were washed to discard unbound neutrophils and fixed for the staining procedure. **e** Images of 3D microfluidic experiments with human neutrophils untreated or stimulated with vimentin, citrullinated vimentin, or LPS (all at 1 µg/mL). Neutrophil elastase (green), actin (red), nuclei (blue). Scale bar, ~200 µm. **f - h** Number of neutrophils that adhere to the HUVECs in the lumen of the microfluidic device (**f**) (n = 3). The number of neutrophils transmigrated through the endothelial monolayer (**g**) (n = 3). Each extravasated neutrophil’s migration distance (**h**) (CT: n =6; Vim: n = 117; CitVim: n = 457; LPS: n = 58). Data are presented as the mean ± standard deviation of the mean. *, P ≤ 0.05; **, P < 0.01; ***, P < 0.001. Significance was determined by one-way ANOVA with Tukey’s test.

### Extracellular vimentin enhances phagocytosis and NETosis

Next, we investigated the impact of eVim and CitVim on neutrophil phagocytosis of fungal and bacterial cells (*C. albicans* and *E. coli, respectively*) (**Fig. 4a**). We either simultaneously introduced eVim or CitVim along with the pathogens to the neutrophils or preincubated the neutrophils with eVim (pre-eVim) or CitVim (pre-CitVim) to assess the requirement for neutrophil priming. Neutrophils alone demonstrated a significant response to both fungal and bacterial pathogens, as evident from the number of neutrophils engulfing these cells. eVim and CitVim increased the number of neutrophils engaged in phagocytosis, except for neutrophils preincubated with CitVim and then exposed to *E. coli* (**Fig. S4**). Further analysis revealed a significant increase in the phagocytosis index (number of ingested pathogens per 100 neutrophils) when eVim or pre-eVim were present, but the effect of CitVim was comparatively lower (**Fig. 4b, d**). Similarly, eVim stimulated more intracellular pathogen killing than CitVim (**Fig. 4 c, e**). We also explored the impact on phagocytosis-related gene expression by exposing neutrophils to eVim or CitVim. eVim triggered more gene expression changes than CitVim (**Fig. 4f**), with these genes being interrelated and mainly associated with phagocytosis (**Fig. S5 and S6**).

**Figure 4.**
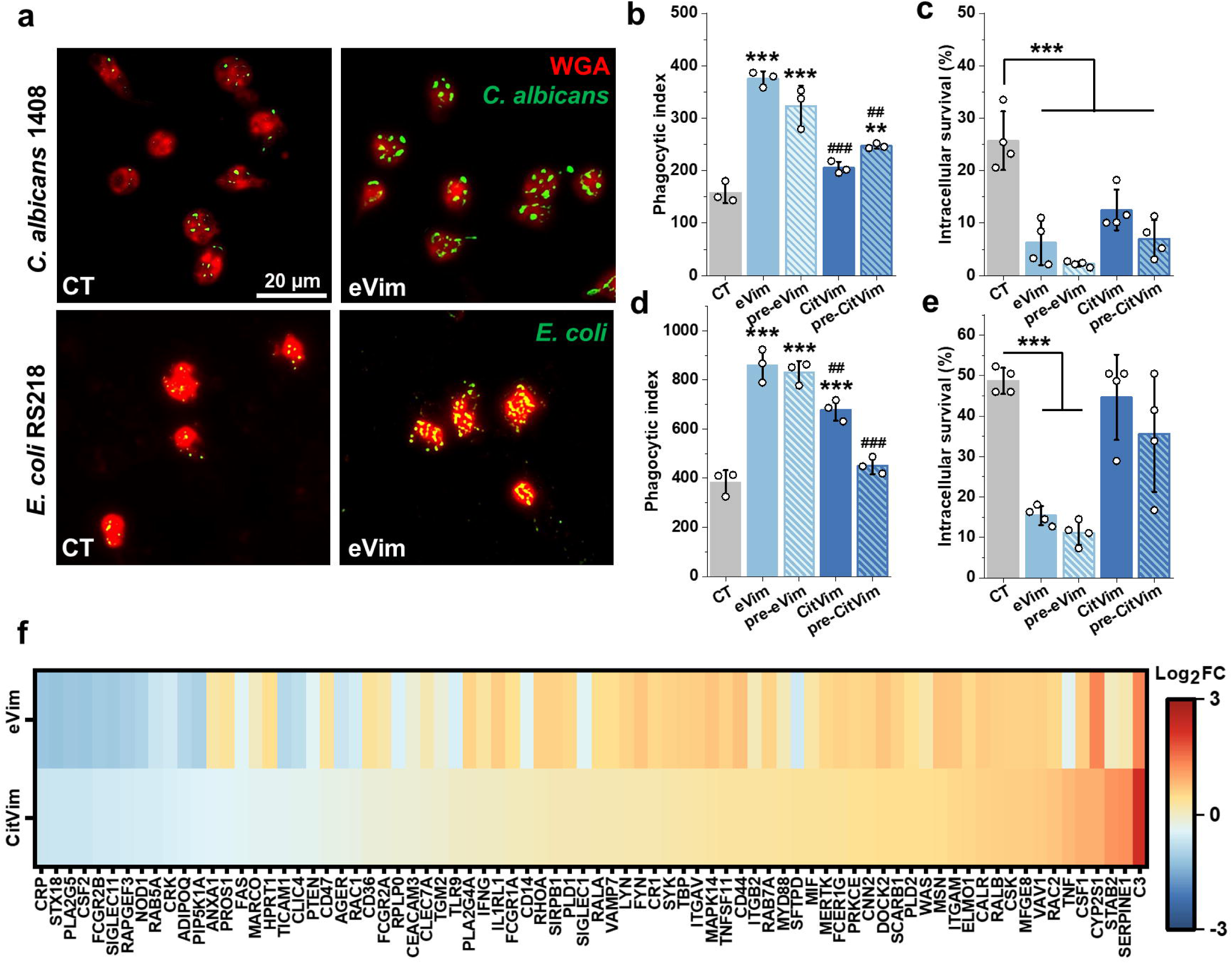
Extracellular vimentin enhances phagocytosis in human neutrophils. **a** Representative images of human neutrophils untreated or treated with vimentin at 1 µg/mL with ingested *Candida albicans* and *Escherichia coli.* Yeast was prestained with Calcofluor White (green pseudocolor), while the bacterium was prestained with Fluorescein isothiocyanate (green). The neutrophil cell membrane was stained with wheat germ agglutinin antibody (WGA – red). Scale bar, ~ 20 µm. **b - g** Phagocytic index for vimentin (eVim – 1 µg/mL) or citrullinated vimentin (CitVim – 1 µg/mL) treated human neutrophils ingesting *C. albicans* cells (n = 3) at MOI of 10 (**b**) and *E. coli* at MOI of 100 (**d**). Additionally, neutrophils were preincubated with Vimentin (pre-eVim) and Citrullinated vimentin (pre-CitVim) at 1 µg/mL for 1 h in cell culture media, then yeasts/bacteria were added to the neutrophils for 2h. The phagocytic index, e.g., the number of fungal and bacterial cells ingested (per 100 cells) by the neutrophils, was manually counted after 2 h of coincubation (n = 3). The intracellular survival of *C. albicans* (**c**) and *E. coli* (**e**) after 2 h infection in human neutrophils. Intracellular survival was evaluated by diluting samples on Sabouraud or McConkey’s agar plates (n = 4). **f** Heat map of changes in gene expression of selected phagocytosis-related genes upon stimulation with vimentin and citrullinated vimentin at 1 µg/mL for 2 h (n = 3). Results are presented as the log_2_ fold change (log_2_FC) compared to the untreated condition (0). The warmer color, the higher gene expression, while the colder color reflects decreased expression. Data are presented as the mean ± standard deviation of the mean. *, P ≤ 0.05; **, ##, P < 0.01; ***, ### P < 0.001. A hash sign (#) indicates a statistical significance comparison between corresponding vimentin and citrullinated vimentin conditions. Significance was determined by one-way ANOVA with Tukey’s test.

To investigate the impact of eVim and its citrullinated form on NET secretion, both types of vimentin were added to neutrophils for 4 hours, and NETosis was assessed by staining for neutrophil elastase (NE) (**Fig. 5a**). Both eVim and CitVim stimulated the release of NETs, but CitVim induced more NETs than eVim (**Fig. 5b**). We also investigated the protein levels and gene expression of citrullinated histone H3 (CitH3), NADPH Oxidase Subunit 2 (NOX2), TLR4, and NE, which are associated with NET formation (**Fig. 5c**). At the protein level, CitVim caused more elevation in NOX2 and NE (**Fig. 5d - g**). At the gene level, CitVim was more effective than eVim in stimulating the expression of all the tested genes associated with NET formation (**Fig. 5h - k**).

**Figure 5.**
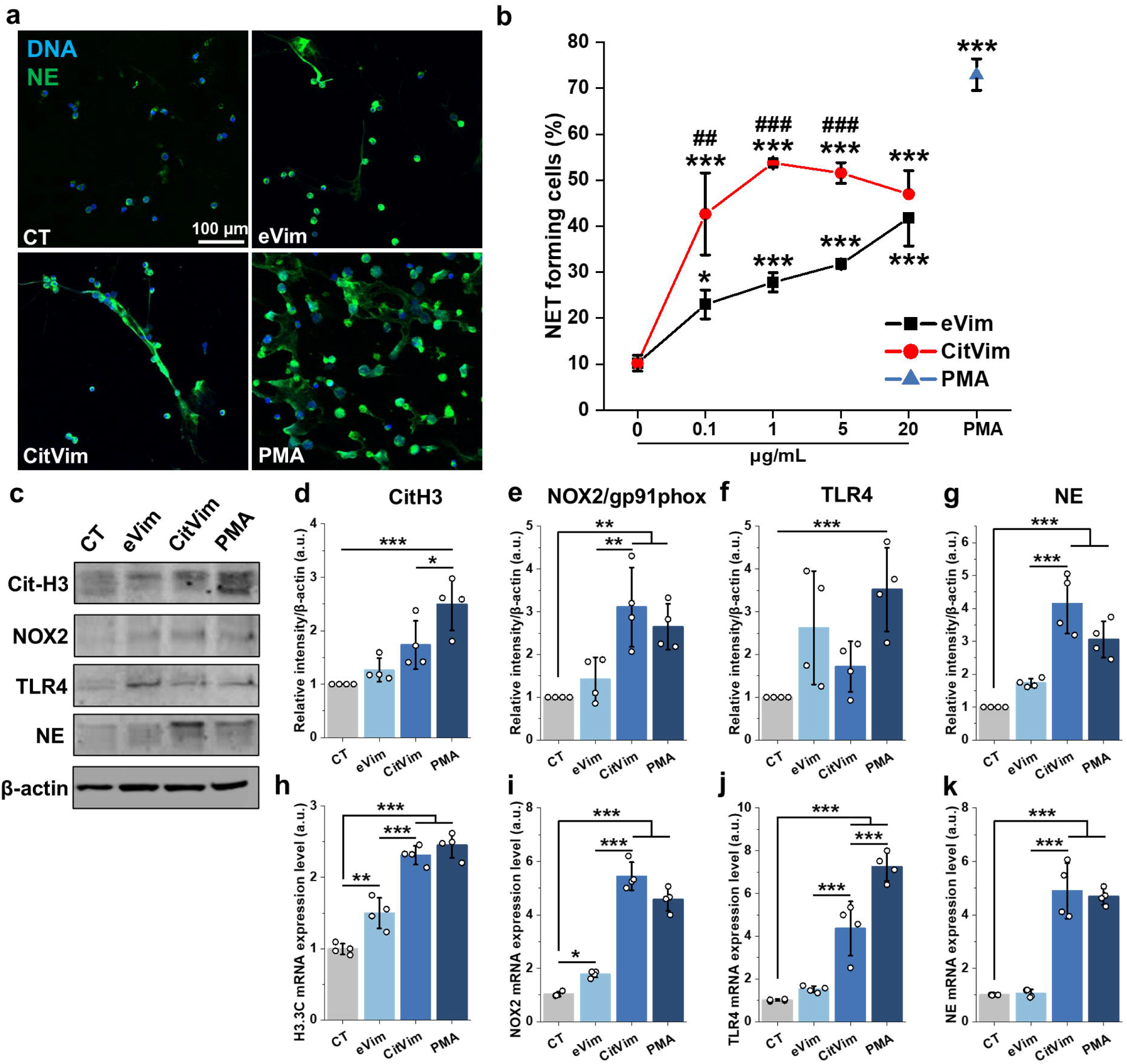
Citrullinated vimentin triggers release of NETs by human neutrophils. **a** Images of NETs released by human neutrophils that were untreated or treated for 4 h with vimentin (1 µg/mL), citrullinated vimentin (1 µg/mL), and PMA (100 nM). Neutrophil elastase (NE - green), nuclei (blue). Scale bar, ~100 μm. **b** Percentage of neutrophil extracellular traps (NETs) released by human neutrophils treated with vimentin (0.1 – 20 µg/mL), citrullinated vimentin (0.1 – 20 µg/mL) or PMA (100 nM) for 4 h (n = 3). **c** Representative immunoblots for the indicated proteins using lysates from human neutrophils that were untreated or treated with vimentin (1 µg/mL), citrullinated vimentin (1 µg/mL), and PMA (100 nM) for 4 h. **d - g** Quantification of the immunoblotting experiments for the protein expression of CitH3 (**d**), NOX2/gp91pox (**e**), TLR4 (**f**), and NE (**g**). Results were normalized to the expression of the β-actin and are presented relative to those of the negative control (CT), set as 1 (n = 3). **h - k** mRNA expression of H3.C3 (**h**), NOX2/gp91phox (**i**), TLR4 (**j**), and NE (**k**). Neutrophils were stimulated with vimentin, citrullinated vimentin, and PMA, as in an immunoblotting experiment. Results were normalized to the GAPDH expression and presented relative to those of the negative control (CT), set as 1. Data are presented as the mean ± standard deviation of the mean. *, P ≤ 0.05; **, P < 0.01; ***, P < 0.001. Significance was determined by one-way ANOVA with Tukey’s test.

Taken together, these results show that eVim, in both its native and citrullinated forms, significantly enhances neutrophil phagocytosis and NETosis. eVim generally promotes phagocytosis more effectively than CitVim, which is especially evident in eVim-preincubated neutrophils. For NETosis, CitVim induces a more robust response than eVim, elevating protein levels and gene expression associated with NET formation to levels comparable to those induced by the potent neutrophil activator (PMA - phorbol 12-myristate 13-acetate) (27).

### Citrullinated vimentin boosts the inflammatory response of human neutrophils, and antibodies targeting vimentin diminish this response

Given the fact that NETs also contain eVim and CitVim, NET formation can potentially result in autocrine and paracrine effects on neutrophils (**Fig. 6a**). We examined the secretion of cytokines and chemokines in response to different doses of eVim and CitVim (**Fig. 6b - i**). CitVim strongly stimulates human neutrophils to produce pro-inflammatory cytokines, specifically IL-1β, IL-6, IL-8, and TNF-α. In contrast, eVim only induced IL-8 secretion. CitVim exhibited a potent effect even at a dose as low as 0.1 µg/mL, with a minimal further increase beyond 1 µg/mL. Additionally, we observed a modest increase (up to 2-fold) in IL-10 secretion following CitVim treatment.

**Figure 6.**
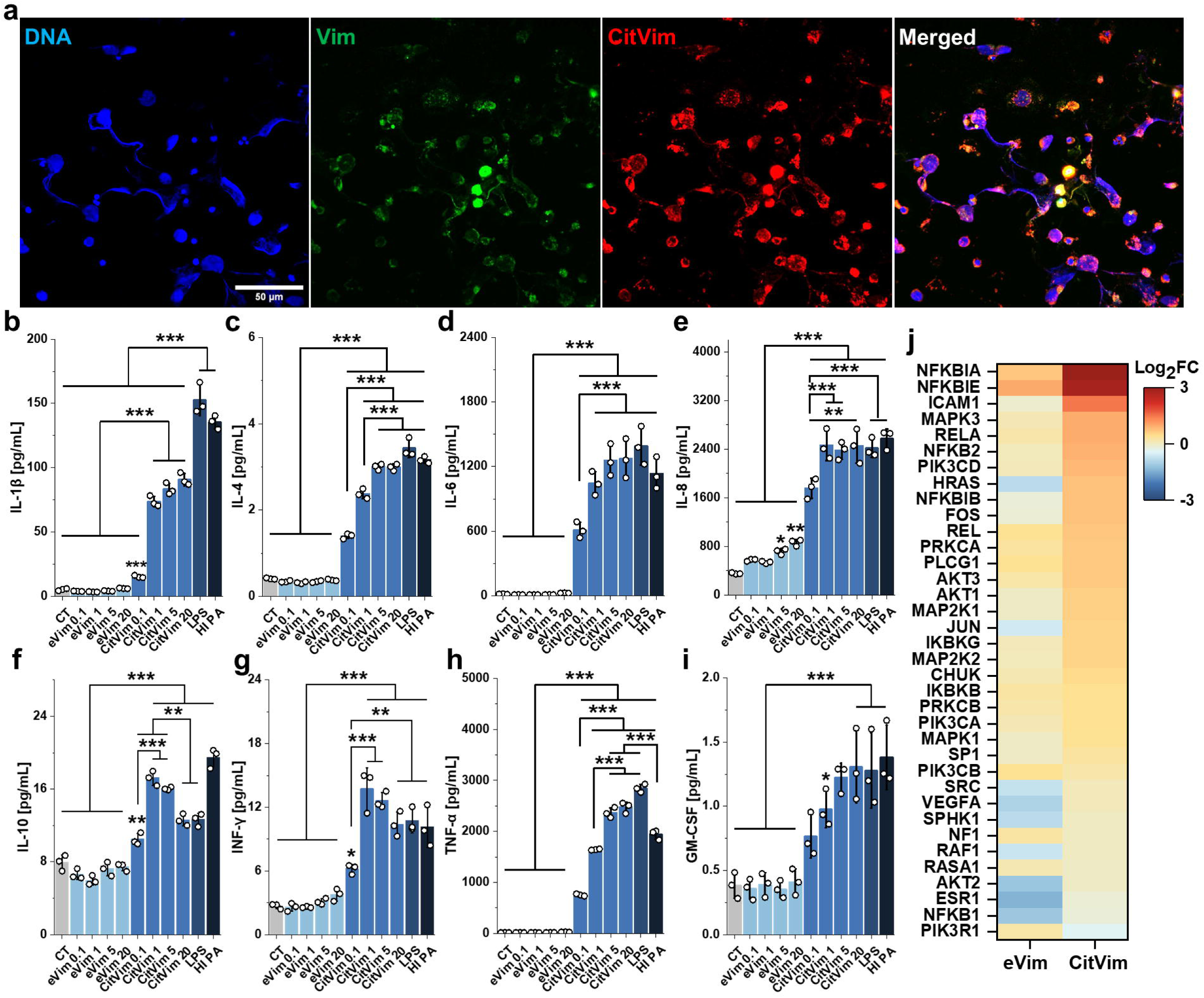
Citrullinated vimentin triggers inflammation in human neutrophils. **a** Representative image indicating the presence of vimentin and citrullinated vimentin within the NETs of PMA (100 nM – 4 h) stimulated neutrophils. Vimentin (green), Citrullinated vimentin (red), DNA (blue). Scale bar, ~50 μm. **b – i** Production of IL-1β (**b**), IL-4 (**c**), IL-6 (**d**), IL-8 (**e**), IL-10 (**f**), IFN-γ (**g**), TNF-α (**h**), and GM-CSF (**i**), as determined by magnetic bead-based enzyme-linked immunosorbent assay (ELISA), in the culture supernatants of human neutrophils that were treated with vimentin (0.1 – 20 µg/mL) and citrullinated vimentin (0.1 – 20 µg/mL). LPS (1 µg/mL) and heat-inactivated *Pseudomonas aeruginosa* (HI PA – 3×10^6^/mL) were used as a positive control (n =3). **j** Heat map of changes in gene expression of selected inflammation-related genes upon stimulation with vimentin and citrullinated vimentin at 1 µg/mL for 4 h (n = 3). Results are presented as the log2 fold change (log2FC) compared to the untreated condition (0). The warmer color, the higher gene expression, while the colder color reflects decreased expression. Data are presented as the mean ± standard deviation of the mean. *, P ≤ 0.05; **; P < 0.01; ***, P < 0.001. Significance was determined by one-way ANOVA with Tukey’s test compared to untreated conditions (CT).

CitVim augmentation of the inflammatory response in human neutrophils is further supported by gene expression analysis, which showed numerous elevated genes following CitVim stimulation, indicating its impact on transcription and inflammation (**Fig. 6j**). These genes are associated with NF-κB activation, consistent with the secretion of pro-inflammatory cytokines. More genes were upregulated by CitVim than by eVim (**Fig. S7 and S8**). These CitVim-triggered genes were found to be strongly co-expressed and were associated with pathways linked to inflammation, including TLR and TNF-α receptor signaling, as well as cancer and viral disease markers (**Fig. S7**).

To assess whether inhibition of CitVim can mitigate cytokine secretion (**Fig. 7a**), various anti-Vim antibodies (**Supplementary Table 1)** were tested for their ability to prevent the secretion of IL-1β, IL-6, IL-8, and TNF-α (**Fig. 7b - e**). Neutrophils were treated with antibodies at varying dilutions (ranging from 1:1000 to 1:100) simultaneously with CitVim at 1 µg/mL for 4 h.

**Figure 7.**
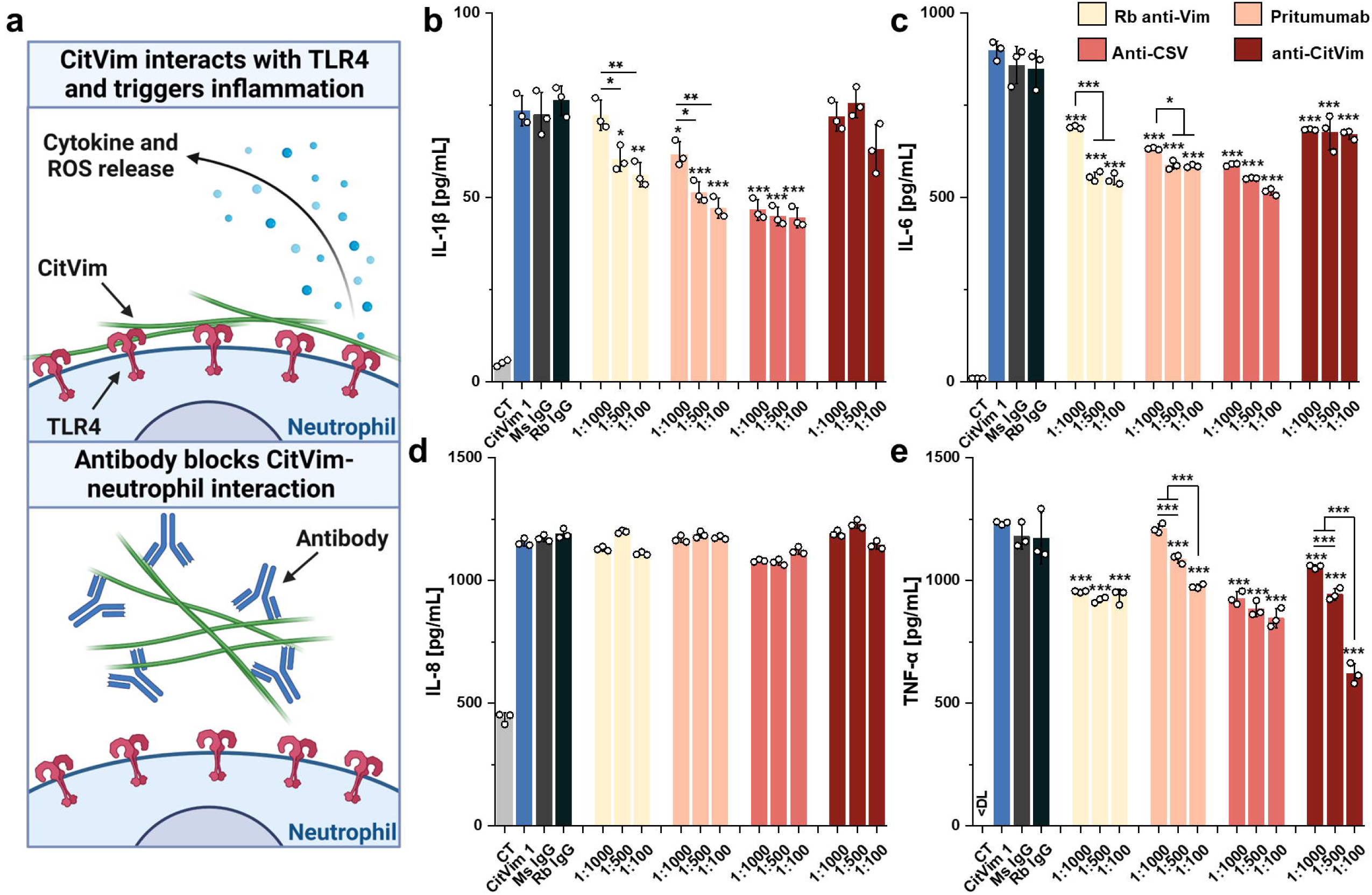
Anti-vimentin antibodies mitigate inflammatory response in human neutrophils caused by citrullinated vimentin. **a** Diagram illustrating the anti-inflammatory effects of anti-vimentin antibodies. These antibodies inhibit the interaction between neutrophil surfaces and free citrullinated vimentin, thereby limiting the inflammatory response. **b – e** Secretion of IL-1β (**b**), IL-6 (**c**), IL-8 (**d**), and TNF-α (**e**) by human neutrophils treated with 1 µg/mL of citrullinated vimentin alone and with the addition of anti-vimentin antibodies (1:1000 – 1:100) (n = 3). Mouse (Ms IgG) and rabbit (Rb IgG) isotype IgG control was used at 0.1 mg/mL. Data are presented as the mean ± standard deviation of the mean. *, P ≤ 0.05; **; P < 0.01; ***, P < 0.001. Significance was determined by one-way ANOVA with Tukey’s test compared to citrullinated vimentin (CitVim 1).

The extent of inhibition of inflammatory mediator secretion depended on the specific antibody used. All antibodies generally had an inhibitory effect on IL-1β, IL-6, and TNF-α secretion. For IL-1β and IL-6, the most pronounced effect was observed with Pritumumab and anti-CSV (cell-surface vimentin) antibodies. In contrast, the secretion of TNF-α was most significantly impacted by the anti-CitVim antibody. None of the antibodies inhibited IL-8 secretion, which might suggest a distinct pathway exploited by CitVim to stimulate IL-8 secretion compared to other mediators.

### In human neutrophils, eVim and CitVim serve as a ligand for Toll-like receptor 4

Our study thus far demonstrates that eVim and CitVim trigger the expression of key immune response regulators: NADPH 2 oxidase, NFκB, and MAPK3 (**Fig. 2e**, **Fig. 5e, i and Fig. 6j**). This activation mirrors the typical upregulation observed in TLR4-dependent immune responses (**Figure 8a**) (28).

**Figure 8.**
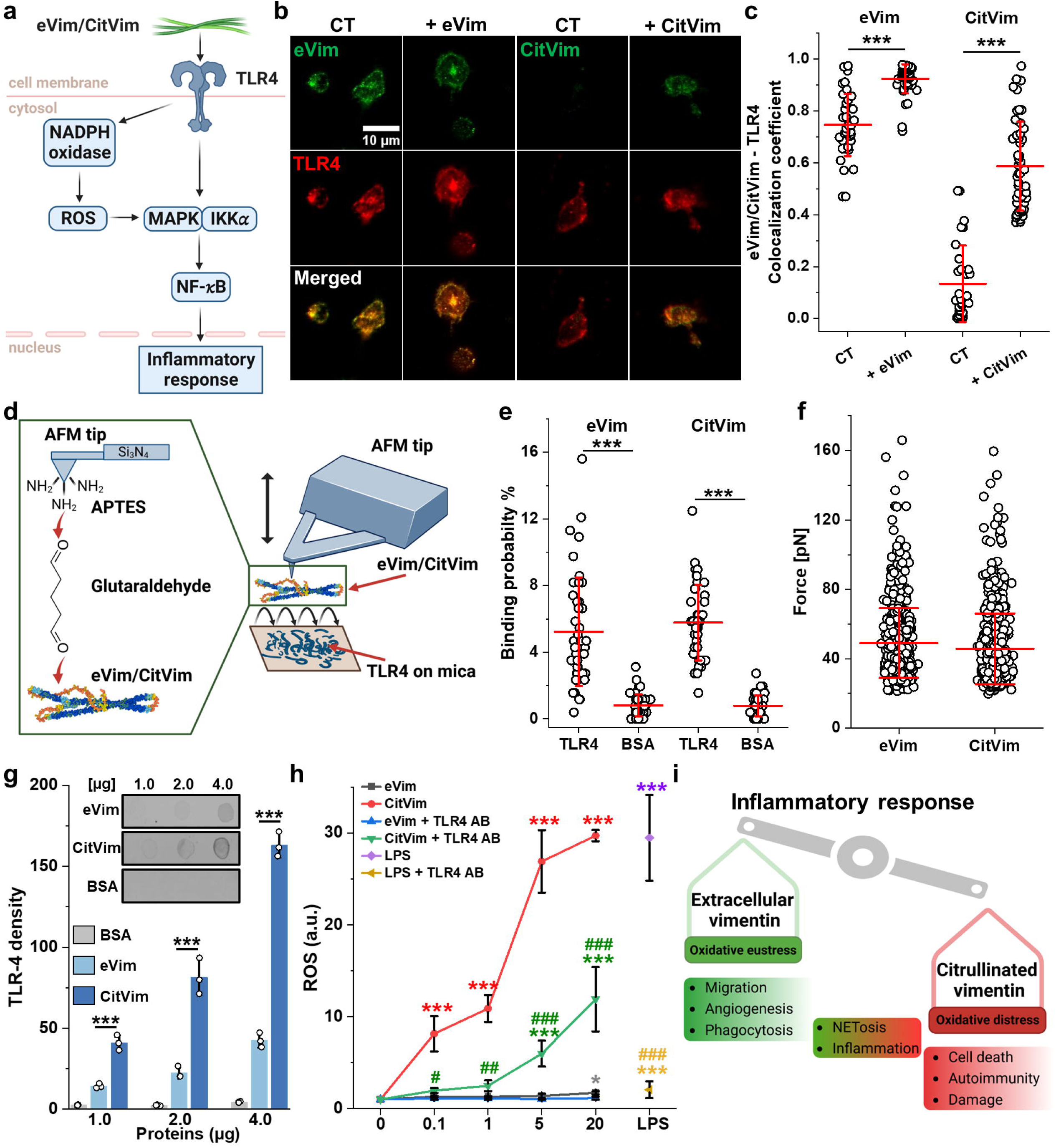
The Toll-like receptor 4 serves as a ligand anchor for eVim and CitVim on the surface of human neutrophils. **a** The presumed mechanism of action of eVim/CitVim on TLR4. **b** Images of human neutrophils showing eVim and CitVim colocalization with TLR4. In control (CT) and treated (+1 µg/mL for 1h) groups, colocalization areas are highlighted in orange. The scale bar, ~10 µm. **c** Quantitative analysis of eVim/CitVim colocalization with TLR4 at the neutrophil membrane. n = 40, 44, 31, and 49 cells for CT (eVim), + eVim at 1 µg/mL, CT (CitVim), and + CitVim (1 µg/mL) groups, respectively. Each symbol represents one cell (n = 3). **d** Functionalization of the AFM tips with eVim and CitVim, for interaction studies. **e** A box plot demonstrates the specific binding probabilities between eVim/CitVim and TLR4-immobilized mica. Each data point on a force map reflects the percentage of adhesive force curves (n = 43). **f** Rupture forces between eVim or CitVim and TLR4 on the mica surface. Each data point corresponds to the rupture force detected for Vim (n = 590) and CitVim (n = 663) derived from 43 force-distance maps. **g** Dot blot assay assessed eVim/CitVim interactions with TLR4, quantified by a bar plot correlating interaction intensities with protein amounts immobilized (n = 3). **h** Reactive oxygen species (ROS) release measured after exposing human neutrophils to varying concentrations of eVim, CitVim, and LPS with and without TLR4 inhibition (TLR-4 antibody). ROS levels were normalized to untreated conditions (n = 3). **i** Diagram showing the imbalance in eVim and CitVim actions relating to inflammatory responses and oxidative stress. Data are presented as the mean ± standard deviation of the mean. *, #, P ≤ 0.05; **, ##, P < 0.01; ***, ### P < 0.001. A hash sign (#) indicates a statistical significance comparison between corresponding eVim/CitVim and LPS compared to conditions with anti-TLR4 antibodies (**h**). Significance was determined by one-way ANOVA with Tukey’s test or Student’s t-test (**f**).

Using confocal microscopy, we next explored the colocalization patterns of TLR4 with eVim and CitVim on neutrophil surfaces (**Fig. 8b, c**). The external application of eVim to neutrophils led to its adsorption and an increased overlap in fluorescence signal, measured by a colocalization coefficient ranging from 0 (no colocalization) to 1 (complete colocalization). Control (CT) neutrophils displayed negligible surface CitVim. However, when CitVim was externally administered, it adhered to the cell surface, exhibiting a colocalization with TLR4 that was less pronounced than with eVim.

To assess the molecular interactions between TLR4 with eVim and CitVim, we used atomic force microscopy (AFM) and force spectroscopy analysis. We affixed either eVim or CitVim to the end of an AFM tip functionalized with glutaraldehyde (**Fig. 8d**). The examination of binding involved recording force-distance (FD) curves through repetitive approaches and retractions of the eVim or CitVim-functionalized tip from a TLR4-coated mica surface as a modification of previous studies (29–31). Specific adhesion events were identified in more than 5% of the retraction FD curves, aligning with findings from other studies on different surface receptor systems (**Fig. 8e**) (29, 30, 32). Control experiments to validate interaction specificity included (i) deploying an AFM tip functionalized only with glutaraldehyde and (ii) directing the tip towards mica surfaces coated with BSA or lacking TLR4 (**Fig. 8e and Fig. S9**). The binding probability observed in these control experiments was notably lower, thus affirming the specificity of the eVim/CitVim and TLR4 complexes under our experimental conditions. The rupture forces detected for the AFM probe at an approach/retraction speed of 5 µm/s were comparable for both eVim and CitVim (**Fig. 8f**), and in the same range as other AFM-based ligand-receptor studies (29, 33).

Dot blot binding assays confirmed the interaction between TLR4 and both eVim and CitVim. Initially, we checked the presence of eVim and CitVim on the nitrocellulose (NC) membrane **(Fig. S10).** Then, NC with immobilized eVim and CitVim was incubated with lysates of human neutrophils to check whether TLR4 would bind to the proteins. BSA served as a negative control. TLR4 binds to both eVim and CitVim but not to BSA (**Fig. 8g**). Density analysis shows that the CitVim – TLR4 interaction occurs at higher affinity than the eVim – TLR4 interaction.

Finally, we employed a functional assay to determine whether blocking TLR4 on the surface of neutrophils inhibits the increased production of ROS caused by eVim and CitVim (**Fig. 8h**). To do so, we used anti-TLR4 antibody at 1:100 and TAK-242, a TLR4 inhibitor at 1 µM, a concentration that did not affect ROS production in untreated conditions (**Fig. S11a**). CitVim caused a significant dose-dependent increase in ROS, which was potently inhibited along with LPS when TLR4 was blocked with anti-TLR4 antibodies. eVim also caused increased ROS formation; however, it was weaker than CitVim. Heat denaturation of eVim or CitVim eliminated their stimulation of ROS production (**Fig. S11b**). The TLR4 inhibitor also partially decreased ROS production induced by CitVim and our positive control with LPS, but did not block ROS production induced by PMA, which acts through a TLR4-independent mechanism (**Fig. S12**). These results demonstrate the specificity of eVim and CitVim to stimulate ROS production by neutrophils and underscore the differential impact of eVim and CitVim on the oxidative balance (**Fig. 8i**).

## Discussion

In this study, we investigated the effects of eVim on human neutrophils to determine the significance of vimentin release to extracellular fluids and the importance of vimentin citrullination for immune responses and possibly immunopathology. We found that CitVim significantly outcompetes its native form (eVim) in the promotion of (i) neutrophil adhesion, (ii) transendothelial migration, (iii) NET formation, (iv) pro-inflammatory response and (v) immune gene expression. In contrast, eVim is more effective than CitVim in inducing neutrophil phagocytic activity. Both forms of vimentin appear to exert these immunomodulatory actions serving as ligands for TLR4.

The presence of Vim on the cell surface results from its release from Vim-expressing cells (11, 21, 34). Our study uncovers a novel binding interaction between Vim and TLR4, adding a crucial dimension to our understanding of Vim’s functional role. This discovery not only broadens the scope of known interactions but also provides insights into novel mechanistic pathways where Vim and TLR4 interplay could be critical in inflammatory and immune responses.

Both eVim and CitVim can be deposited on the cell surface. However, the distribution of CitVim differs from eVim, possibly due to differences in the binding properties of both vimentins with other cell surface components. TLR4 is unlikely to be the only attachment point for Vim on the neutrophil surface, since previous studies have shown binding of eVim to N-acetylglucosamine, glycosaminoglycans, and several transmembrane proteins (23, 35–37). On hydrogels coated with eVim and CitVim, both vimentins promote neutrophil adhesion, clustering, and aggregation. This phenomenon is consistent with previous studies demonstrating that neutrophils tend to aggregate in response to external stimulation with inflammatory mediators such as granulocyte-macrophage colony-stimulating factor (GM-CSF) or PMA, mediated by the β2-integrin ligand CD54 (38). Furthermore, two independent groups have demonstrated that ICAM-1 plays a pivotal role in neutrophil activation and aggregation, highlighting its involvement in increased adherence and aggregation upon exposure to inflammatory factors also in a murine model of endotoxemia (39, 40).

The capacity of eVim to enhance transendothelial migration suggests that eVim released from damaged or stressed cells into the extracellular space can influence the extent of neutrophil infiltration into tissues and could impact the degree of tissue damage. Our results suggest that both eVim and CitVim act as DAMPs in the modulation of the immune response, particularly by activation of neutrophils. In this light, the release of NETs containing CitVim during neutrophil activation might act as a positive feedback loop that propagates neutrophil activation within sites of inflammation.

Elevated eVim levels are prevalent in severe conditions like sepsis, idiopathic pulmonary fibrosis (IPF), and viral infections, with Intensive Care Unit (ICU) patients showing average serum Vim concentrations around 500 ng/mL with levels nearly triple this amount observed in some patients (14, 41–43). Notably, during inflammation, eVim and CitVim levels at the site often surpass peripheral blood levels, as evidenced by findings of higher CitVim concentrations in lung tissue and bronchoalveolar lavage fluid of IPF patients compared to their bloodstream (14). Moreover, reported eVim concentration in COVID-19 patient saliva can reach up to 70 µg/mL (44). These findings underline the relevance and necessity of the diverse range of eVim and CitVim doses (0.1 - 20 µg/mL) employed in our study, reflecting the complex and varying nature of eVim and CitVim levels in different disease states.

The current understanding of eVim’s role in the host inflammatory response includes several different cellular or molecular effects and different conclusions about whether eVim systemically is pro- or anti-inflammatory. For example, eVim modulates oxLDL-induced secretion of pro-inflammatory cytokines in macrophages and alters cytokine profiles in LPS-activated dendritic cells, affecting T cell differentiation (45, 46). Additionally, its involvement in diapedesis, influencing the distribution and expression of adhesion molecules in lymphocytes and endothelial cells, highlights its multifaceted role (47). eVim’s stimulatory effect on leukemic macrophage migration and phagocytosis aligns with our findings on human neutrophils (48). Secretion of vimentin by tumor endothelial cells can lead to immunosuppression, and vaccination to produce anti-vimentin antibodies shows efficacy in prolonging survival in dogs with bladder cancer (49). eVim’s capability as a substrate for attachment and motility of various cell types underscores its diverse, context-dependent interactions (23). Collectively, these findings highlight the pivotal role of eVim in orchestrating key elements of the immune response, revealing its potential as a critical target in therapeutic interventions aimed at modulating inflammatory processes.

One aspect of eVim that might determine its effect *in vivo* is the state of vimentin citrullination. Cytoskeletal vimentin needs to be disassembled before it can be secreted, and this disassembly appears to involve either citrullination or phosphorylation. In neutrophils, citrullination appears to be required, whereas, in macrophages, phosphorylation appears to be dominant (19, 50, 51). It is not yet clear whether the effect of citrullinating vimentin is to reduce the filaments to small soluble oligomers, or whether the change from arginine to citrulline creates new binding sites on the cell or ECM. The impact of citrullination on protein function is complex and varies significantly. For example, citrullination diminishes the osteoprotective effect of fibrinogen, suggesting a detrimental impact in this context (52). In contrast, citrullination of histone H4 leads to a decrease in calcium influx compared to its native form, resulting in reduced NET formation (53). Citrullinated collagen, as opposed to its native form, does not decrease the severity of autoimmune arthritis (54). These examples highlight that citrullination can lead to diverse functional outcomes, ranging from loss of beneficial properties to a reduction in potentially harmful activities, depending on the specific protein undergoing modification.

CitVim specifically binds to the human leukocyte antigen class II histocompatibility antigen, DRB1 beta chain (HLA-DRB1) in rheumatoid arthritis (RA) patients, enhancing cytokine production, notably INF-γ (55). CitVim increases proliferation and cytokine secretion in RA and osteoarthritis patient-derived fibroblast-like synoviocytes (FLS), contrasting with the inhibitory effects of non-modified Vim (56). Additionally, CitVim induces TGF-β1, CTGF, and IL-8 secretion in lung fibroblasts and increases bone resorption in a mouse periodontitis model (14, 57). Our research aligns with these findings, revealing that vimentin-targeting antibodies inhibit CitVim-mediated pro-inflammatory cytokine secretion but not IL-8. This is partially corroborated by data showing that NF-κB inhibition reduces specific cytokines in RA FLS, suggesting that anti-vimentin antibodies may specifically inhibit NF-κB pathways (58).

## Conclusions

Overall, our study shows the efficiency of eVim to activate neutrophils and suggests that this activation can lead to extravasation and augmentation of inflammatory processes. The data also highlight the role of TLR4 in the differential response of neutrophils to eVim and CitVim. CitVim exhibits enhanced activation of some neutrophil functions, whereas it is less effective than eVim in others, and antibodies targeting Vim can mitigate the inflammatory response caused by CitVim. These findings offer insights into potential strategies for manipulating vimentin-related processes to control inflammation and reduce tissue damage in inflammatory disorders.

## Materials and Methods

### Neutrophil isolation

Blood was collected from healthy donors with the approval of the Bioethics Committee at the Medical University of Bialystok (APK.002.8.2023). Neutrophils were isolated by density gradient centrifugation using PolymorphPrep (Progen, Heidelberg, Germany). Cells were counted on a hemocytometer and suspended in Dulbecco’s Modified Eagle’s Medium (DMEM) supplemented with 10 % FBS (#P30-8500, PAN Biotech) and 1% antibiotic antimycotic solution (#A5955, Sigma-Aldrich, St. Louis, MO, USA).

### Cell culture and treatment

Human umbilical vein endothelial cells (HUVECs, #CRL-1730™, Sigma-Aldrich) were maintained in Endothelial Cell Growth Medium (#211-500, San Diego, CA, USA). Wild-type mouse embryonic fibroblasts (vim +/+ mEFs) and their variant lacking vimentin expression (vim−/− mEFs) were grown in DMEM supplemented with 10 % FBS and 1 % antibiotic antimycotic solution. Cells were grown in a humidified atmosphere with 5 % CO_2_ at 37 °C.

For the experiments, cells were treated with recombinant human vimentin (#10028-H08B, SinoBiological, Wayne, PA, USA) and recombinant human citrullinated vimentin (#21942, Cayman Chemicals, Ellsworth, MI, USA), both at a concentration range of 0.1 to 20 µg/mL. Lipopolysaccharide from *Escherichia coli* O26:B6 (LPS, 1 µg/mL, #L8274, Sigma-Aldrich) and phorbol 12-myristate 13-acetate (PMA, 100 nM, #10008014, Cayman Chemicals) were used as positive controls.

### Vimentin staining

To evaluate whether Vim and CitVim can bind the surface of cells, mEFs −/− and HUVECs were grown on a glass coverslip. When cells reached the desired confluence, Vim and CitVim (both at 1 µg/mL) were introduced or not (CT) to the cells for 1 h. Then, the unbound protein was washed 3 times with PBS, and cells were fixed in 3.7 % paraformaldehyde (PFA) for 30 min at RT. Next, cells were permeabilized with 0.1 % Triton X-100 for 15 min at RT, followed by 30 min incubation with 0.1 % bovine serum albumin (BSA) to block nonspecific binding of the secondary antibody. Cells were stained with rabbit anti-vimentin (#ab45939, Abcam, Cambridge, UK) or mouse anti-citrullinated vimentin (#22054, Cayman Chemicals) at a dilution of 1:1,000 for, RT. The cells were further stained with a secondary anti-rabbit (#ab150081, Abcam) and anti-mouse (#ab150113, Abcam) antibody conjugated with Alexa Fluor 488 dye at a dilution of 1:1,000 for 1 h at RT in the dark. Actin was stained with Texas Red™-X Phalloidin (#T7471, Invitrogen, Waltham, MA, USA). The cell nuclei were counterstained with Hoechst 33342 (#R37605, Invitrogen). The coverslips were mounted with an antifade fluorescence mounting medium (Abcam) and examined by confocal microscopy.

Since endothelial cells express intracellular vimentin, vital staining was performed to visualize extracellular vimentin. To do so, anti-Vim antibodies (at 1:500) were introduced to the non-fixed cells for 1 h. Cells were incubated with antibodies at 4 °C to minimize possible endocytosis of the antibodies. Then, the antibodies were washed thrice with PBS, and the cells were fixed. After fixation, vital stained cells were permeabilized and stained with secondary antibody, Texas Red™-X phalloidin and Hoechst 33342, to visualize eVim, actin, and nuclei, respectively.

### Adhesion assay

To assess if the presence of extracellular eVim/CitVim on the surface of the cells determines the adhesion of the immune cells, vim−/− mEFs and mEF +/+ were grown in 96-well black cell culture plates with optical bottom until full confluency was reached. Then, vim−/− mEFs were preincubated with eVim and CitVim (both at the range of 0.1 – 20 µg/mL) for 1 h, and cells were washed three times with PBS. Then, 3 × 10^5^/well of neutrophils were introduced to the confluent monolayer of fibroblasts for 1 h. After 1 h incubation, unbounded neutrophils were washed off with PBS. Cells were fixed with 3.7 % PFA, permeabilized using 0.1 % Triton X-100, and blocked with 0.1 % BSA. Next, the cells were stained with mouse anti-myeloperoxidase (MPO) antibody (1:500, #MA1-80878, Sigma-Aldrich) for 1 h at RT. The cells were stained with a secondary anti-mouse antibody conjugated with Alexa Fluor 647 dye (1:1,000, #ab150115, Abcam) for 1 h at RT in the dark. Actin was stained with AlexaFluor488-phalloidin (#R37110, Invitrogen), and nuclei were stained with Hoechst 33342.

In a parallel experiment, neutrophils were live-stained with calcein AM (1 µM, #C1430, Invitrogen) for 30 min in the dark. Then, neutrophils were washed with PBS by gentle centrifugation. Next, 3 × 10^5^/well of neutrophils were introduced to the confluent monolayer of fibroblasts for 1 h. vim−/− mEFs cells were preincubated with eVim or CitVim as before for 1 h. Unbounded neutrophils were washed off with PBS. The fluorescence (494/517 nm) from calcein AM stained neutrophils was recorded on the Varioskan Lux microplate reader (Thermo Fisher Scientific). The background fluorescence was determined for each condition and subtracted from the total fluorescence values before data analysis. The results were compared to the untreated control, normalized to 1.0, and presented as a fold change in the adhesion of the neutrophils. LPS (1 µg/mL) was added to the neutrophils as a positive control.

### Polyacrylamide hydrogels

Polyacrylamide hydrogels with a stiffness of 30 kPa were prepared using previously described methods (59). 40 % acrylamide (1170 µL, #1610140, Bio-rad) and 2 % bis-acrylamide solutions (594 µL, #1610142, Bio-rad) were formulated in distilled water (2736 µL) to a total volume of 4.5 mL, divided into 500 µL aliquots. To initiate polymerization, 1.5 μL of TEMED (#1610800, Bio-rad) and 3 μL of 10 % (w/v) ammonium persulfate (#1610700, Bio-rad) were added to 500 μL aliquots. Immediately after the addition of polymerization initiators, 190 µl of the solution was pipetted to a 20-mm round glass coverslip (adhesive) pretreated with 5% 3-aminopropyltrimethoxysilane and 0.5% (v/v), glutaraldehyde. The solution was covered with a 24×60 mm glass coverslip (non-adhesive) siliconized with 10 % SurfaSil solution (#TS42800, Thermo Fisher). After 30 min, the top coverslip was removed, and hydrogels were covalently linked to ligands by incubating the gels for 1 h with 50 μL of collagen I, vimentin, or citrullinated vimentin (all proteins at 0.1 mg/mL in 50 mM HEPES pH 8.5) after activating the gel surface with the UV-sensitive Sulfo-SANPAH cross-linker. Before experiments, hydrogels were sterilized in UV light and preincubated with a complete cell culture medium containing antibiotics. The stiffness of the gel substrates was verified using a rheometer (data not shown).

To assess the adhesion and activation of human neutrophils on the surface of 30 kPa polyacrylamide hydrogels coated with collagen, vimentin, or citrullinated vimentin, 2×10^5^ of neutrophils in 200 µL of complete DMEM were added on top of the hydrogels for 1 h. Then, unbounded neutrophils were washed off with PBS. Bright-field images of neutrophils remaining on hydrogels were captured using an inverted microscope. Cell morphology, number of adherent neutrophils, and aggregation index were further processed from the images using ImageJ Fiji Software. The aggregation index is the number of neutrophils within the cluster (at least 3 cells stick together) per 100 cells. After bright-field image acquisition, cells on top of hydrogels were fixed, permeabilized, and blocked. The cells on hydrogels were stained with rabbit anti-NOX2/gp91phox (1:200, #BS-3889R, Invitrogen) for 1 h, at RT. The cells were stained with a secondary anti-mouse antibody conjugated with Alexa Fluor 647 (1:1,000, #ab150115, Abcam) and a secondary anti-rabbit conjugated with Alexa Fluor 488 (1:1,000, #ab150081, Abcam) for 1 h, at RT in the dark. Nuclei were stained as before with Hoechst 33342. The fluorescence intensity of NOX2/gp91phox was determined in ImageJ Fiji software and processed according to the equation:

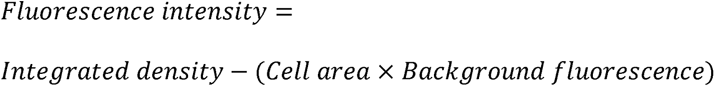

The results were compared to the values in collagen control and presented as a fold change in NOX2 intensity.

### 2D migration assay on Transwell inserts

The migration of neutrophils was assessed on Transwell inserts with a 5 µm pore size semi-permeable membrane (#3421, Corning, NY, USA). The membrane was incubated overnight with 0.1 mg/mL of collagen, vimentin, or citrullinated vimentin at 37 °C. The membrane was washed with PBS. 3×10^5^ of neutrophils in 200 µL of the medium was added for 2 h to the upper chamber of the Transwell insert, and the bottom chamber was filled with 500 µL of media. After 2 h, media from the bottom chamber was collected, and the number of neutrophils that migrated through the membrane was counted on a flow cytometer using Muse Count & Viability Kit (#LUMIMCH100102, Merck, NJ, USA) according to the producer’s instructions. The upper chamber was washed with PBS thrice, fixed in 3.7 % PFA, stained with Hoechst 33342, and imaged on a fluorescence microscope.

### Microfabrication of the 3D microfluidic device

We followed a previously described method with some modifications to fabricate the microfluidic device (60). Initially, a 3-inch silicon wafer was prepared, and the SU-8 2025 epoxy-based photoresist (MicroChem) was carefully poured onto it at 95 °C. The wafer with the photoresist was then left overnight for curing. Next, a photomask was precisely aligned and applied over the wafer, which was then exposed to a 200-W UV lamp for 2 hours. This exposure ensured the polymerization of the photoresist, forming the desired microstructures. After exposure, any unpolymerized photoresist was dissolved using propylene glycol methyl ether acetate (PGMEA) (MicroChem). Polydimethylsiloxane (PDMS) (#2065622, Ellsworth) was used to create negative molds for the microfluidic channels. PDMS was poured over the silicon master, replicating the microstructures present on the wafer. Subsequently, positive stamps made of PDMS were cast using the same procedure, generating the microfluidic channels on glass coverslips. Before proceeding with gel fabrication, the hydrogel reservoir of the device was prepared. It was filled with 5 M sulfuric acid (#258105, Sigma-Aldrich) and left for 90 minutes. Afterward, thorough washing with distilled water was performed to remove residual acid. Finally, the reservoir was filled with a solution containing 50 µg/mL collagen type I, and sterilization was achieved using UV light exposure.

### 3D transendothelial migration assay

As previously described (61), each microfluidic device was filled with 60 µL of a hydrogel formulation containing the following components: 6 µL of 10x phosphate-buffered saline (PBS) (#5493, Sigma), 6 µL of 0.1 M sodium hydroxide (NaOH) (#BA0981118, POCh SA), 18 µL of distilled water, and 30 µL of a 10 mg/mL collagen type I solution. This formulation resulted in a gel with a final 5 mg/mL collagen concentration. All components were kept on ice until ready for mixing and then injected into each device’s hydrogel reservoir. To create cylindrical voids resembling human vessels, two freshly coated with sterile 1% BSA 180-mm-diameter acupuncture needles were inserted into the needle guides of the device. The devices were then transferred to a 37 °C environment for 10 minutes to facilitate gel polymerization. To prevent drying of the hydrogel, phosphate-buffered saline (PBS) was pipetted onto the reservoir ports. After 2 hours of incubation, the acupuncture needles were carefully removed from the hydrogel, forming cylindrical voids. Human umbilical vein endothelial cells (HUVECs) were injected into one of the channels at 10 million cells per mL density and left to incubate for 10 minutes. Subsequently, the devices were inverted for an additional 10 minutes to ensure complete coverage of the cylindrical voids by the HUVECs. The cell-seeded devices were then placed in wells of 6-well culture plates containing 7 mL of complete growth medium and incubated in a cell culture incubator for 24 hours. Once the HUVEC monolayer had fully covered the cylindrical voids after 24 hours, 20 µL of human neutrophils at a concentration of 1×10^6^ cells/mL in medium, with or without vimentin, citrullinated vimentin or lipopolysaccharide (LPS) (all at 1 µg/mL), were introduced into the freshly washed lumen of endothelial cells and incubated for 4 hours. Following the 4-hour incubation, the channels were gently washed with media and then fixed with 3.7% paraformaldehyde (PFA) for immunofluorescence staining. After fixation, the hydrogels were carefully detached from the device hydrogel reservoir and placed in 1.5 mL centrifuge tubes. To permeabilize the cells within the hydrogel, 0.2% Triton X-100 was added and incubated for 30 minutes at room temperature. Subsequently, the hydrogels were blocked with 3% BSA for 30 minutes at 37 °C. The hydrogels were then incubated overnight at 4 °C with a rabbit anti-neutrophil elastase antibody (#ab21595, Abcam). After thorough washing, the hydrogels were incubated with the appropriate secondary antibody at a dilution of 1:1000, along with Texas Red™-X Phalloidin and Hoechst 33342, as previously mentioned. Finally, the hydrogels were imaged using a scanning confocal system.

### Fungal and bacterial preparation

Laboratory strains of *Candida albicans* (*C. albicans* 1408) were obtained from the Polish Collection of Microorganisms, Polish Academy of Science in Wroclaw, Poland. The *C. albicans* strain was plated on Sabouraud dextrose with chloramphenicol agar (#PS192, Biomaxima, Lublin, Poland). The clinical strain of *Escherichia coli* (*E. coli* RS218) was plated on MacConkey agar plates (#PS10, Biomaxima). Both strains were grown routinely at 37 °C. Hemocytometer counts were used to estimate the quantity of yeast and bacteria. For coculture with human neutrophils, exponentially growing cells were suspended and diluted to the desired cell number in cell culture media without antibiotics and antimycotics.

### Phagocytosis assays

*C. albicans* 1408 was stained with calcofluor white (#F3543, Sigma-Aldrich) at 0.25 µg/mL for 30 min, RT. *E. coli* RS218 was stained with Fluorescein isothiocyanate (FITC, #F7250, Sigma-Aldrich). Briefly, bacterial cells were resuspended in 0.1 M sodium bicarbonate buffer, pH 9.0. Freshly made FITC (2 µg/mL final concentration) was added to the bacterial suspension for 30 min, RT, in the dark. Then, cells were washed by centrifugation thrice in PBS. Both yeast and bacteria were finally resuspended in cell culture media at a desired concentration.

Prestained, *C. albicans* cells at MOI (multiplicity of infection, number of germ cells per single neutrophil) of 10 and *E. coli* at MOI of 100 were added for 2 h to 1×10*^5^* neutrophils seeded onto 0.01% poly-l-lysine (Sigma-Aldrich) treated 96-well black cell culture plates with the optical bottom. Neutrophils were treated simultaneously with yeast or bacteria and vimentin (eVim – 1 µg/mL) or citrullinated vimentin (CitVim – 1 µg/mL). Alternatively, neutrophils were preincubated with Vimentin (pre-eVim) and Citrullinated vimentin (pre-CitVim) at 1 µg/mL for 1 h in cell culture media. Proteins were washed off, and *C. albicans and E. coli* were added to the neutrophils for 2h. After 2 h of incubation, neutrophils were washed with PBS, fixed with 3.7 % PFA, and blocked with 0.1 % BSA. Neutrophils were stained with Wheat Germ Agglutinin (WGA at 5 µg/mL) conjugated with Alexa Fluor 647 (#W32466, Invitrogen) for 10 min at RT. After staining, cells were examined by fluorescence microscopy. The results are presented as the percentage of cells taking up or adherent to fungal/bacterial cells (engaged neutrophils) and the total number of fungal/bacterial cells taken up per 100 cells (phagocytic index). Data were obtained from 3 separate experiments by analyzing 5 to 10 individual randomly taken images per replicate (at least 500 neutrophils/replicate).

### Intracellular survival assay

Neutrophils (1×10^6^ cells/well in a 24-well plate) were treated as in the phagocytosis assay. After 2 h of incubation with yeast or bacteria, the cultures were collected using cell scrapers and transferred to Eppendorf tubes. The neutrophils were lysed by sonication for 10 min. Serial dilutions were plated on Sabouraud or MacConkey agar for yeast/bacterial outgrowth assessment and left for 24 h of incubation at 37 °C.

### NETosis assay

To assess the formation of neutrophil extracellular traps (NETs), 1 × 10^5^ cells were seeded onto 0.01% poly-l-lysine functionalized sterilized glass coverslips. Neutrophils were untreated or treated with eVim (0.1 – 20 µg/mL), CitVim (0.1 – 20 µg/mL), or PMA (100 nM) for 4 h at 37 °C and 5% CO_2_. After incubation, the cells were washed with PBS, fixed in 3.7% PFA, permeabilized with 0.1% Triton X-100, and blocked in 0.1% bovine serum albumin, as mentioned above. Next, the cells were stained with rabbit anti-neutrophil elastase (NE) antibody (#ab21595; Abcam) at a dilution of 1:500 for 1 h, at RT. The cells were stained with an appropriate secondary anti-rabbit antibody at a dilution of 1:1,000 for 1 h, at RT in the dark. The cell nuclei were counterstained with Hoechst 33342. The coverslips were mounted with an antifade fluorescence mounting medium (#ab104135, Abcam) and examined by confocal microscopy. The number of cells producing NETs was manually counted. At least five images per coverslip were taken randomly and analyzed (n = 3).

### RNA isolation and gene expression

Total RNA was extracted using the universal RNA purification kit (#E3598-02; EURx, Gdansk, Poland) from 5 × 10^6^ neutrophils per well seeded in 6-well cell culture plates. Cells were untreated or treated with eVim (0.1 – 20 µg/mL), CitVim (0.1 – 20 µg/mL), or PMA (100 nM) for 2 h (for phagocytosis-related genes) or 4 h (for NET- and Inflammation-related genes) at 37 °C and 5% CO_2_. The cells were scratched, transferred to Eppendorf tubes, and centrifuged, and the supernatant was discarded. The concentration and purity of the isolated RNA were evaluated using a Qubit 4 fluorometer (Thermo Fisher Scientific). cDNA was synthesized using the iScript cDNA synthesis kit (#1708891; Bio-Rad). Reverse transcription-quantitative PCR (qRT-PCR) was performed with 100 ng of cDNA in a 20-μL reaction mixture containing SsoAdvanced universal SYBR green supermix (#1725274; Bio-Rad) using phagocytosis PrimePCR plates (#10047255; Bio-Rad) and VEGF signaling and activation PrimePCR plates (#10025756, Bio-Rad) on the CFX Opus real-time PCR detection system (Bio-Rad) with the following amplification program: 2 min at 95°C, followed by 40 cycles of 5 s at 95°C and 30 s at 60°C. GAPDH (glyceraldehyde-3-phosphate dehydrogenase) was used as an internal control. The gene expression levels were reported as the relative quantity, expressed using the comparative cycle threshold (CT) method (2^-ΔΔCt^), and presented as log_2_FC.

For quantifying NETosis-related mRNAs, the following primer pairs were used:

*H3.3C*:

Forward 5’-TCCAGAGGTTGGTGAGGGAGAT-3’

Reverse 5’-TAGCGTGGATGGCACACAGGTT-3’

*NOX2*:

Forward 5’-CAAGATGCGTGGAAACTACCTAAGAT-3’

Reverse 5’-TCCCTGCTCCCACTAACATCA-3’

*TLR4*:

Forward 5’-CCCTGAGGCATTTAGGCAGCTA-3’

Reverse 5’-AGGTAGAGAGGTGGCTTAGGCT-3’

*NE*:

Forward 5’-TCCACGGAATTGCCTCCTTC-3’

Reverse 5’-CCTCGGAGCGTTGGATGATA-3’

*GAPDH*:

Forward 5’-GTCTCCTCTGACTTCAACAGCG-3’

Reverse 5’-ACCACCCTGTTGCTGTAGCCAA-3’

### Protein preparation and western blotting

5×10^6^ of human neutrophils were seeded in 6-well plates and were untreated or treated with eVim (1 µg/mL), CitVim (1 µg/mL), or PMA (100 nM) for 4 h. After 4 h, the supernatant was aspirated, cells detached using scrappers, transferred to Eppendorf tubes containing supernatant, and centrifuged. The supernatant was discarded, and the whole-cell lysate was prepared using RIPA lysis buffer (#89901, ThermoFisher) with Pierce Protease Inhibitor (#A32963, ThermoFisher) added freshly before use. Cells were lysed for 20 min on ice and centrifuged at 14,000 rpm for 20 min at 4 °C. Next, supernatants were transferred to fresh tubes, and the Bradford (#5000006, Bio-Rad) assay was performed to determine protein concentration. Lysates were subjected to electrophoresis using 10% sodium dodecyl sulfate–polyacrylamide (SDS-PAGE) at an amount of 15 µg protein per lane. After SDS-PAGE separation, proteins were blotted onto methanol-activated polyvinylidene fluoride (PVDF) membranes. Next, the membranes were blocked for 1 h in 5% nonfat dry milk in TBS-T (150 mM NaCl, 50 mM Tris-base, 0.05% Tween 20, pH = 7.4). Blocked protein blots were incubated with anti-TLR4 (1:1,000; #ab13556, Abcam), anti-NOX2/gp91phox (1:200, #BS-3889R, Invitrogen), anti-neutrophil elastase (1:1,000; #ab21595, Abcam), anti-Cit H3 (1:1,000; #ab10799, Abcam), and anti-β-actin (1:5,000; #A5441, Sigma-Aldrich), in TBS-T at 4 °C overnight, followed by incubation with goat anti-rabbit IRDye 800CW IgG and goat anti-mouse IRDye 800CW IgG secondary antibody in TBS-T (1:10,000) at room temperature for 1 h, in the dark. Protein blots were visualized with the Odyssey LiCor Imaging System (LiCor Biosciences). Band intensities were quantified using Image Studio Acquisition Software. Data are presented as relative intensity of protein of interest bands in eVim, CitVim, or PMA-treated samples compared to the untreated samples and normalized β-actin.

### Cytokine and chemokine secretion

The Bio-Plex Pro human cytokine assay assessed the secretion of IL-4, IL-6, IL-8, IL-10, IFN-γ, TNF-α, and GM-CSF (Bio-Rad Laboratories, Hercules, USA). IL-1β was evaluated using an IL-1 beta Human ELISA Kit (Invitrogen). Neutrophils (3 × 10^5^/well) were cultured on 96-well culture plates for 4 h with vimentin and citrullinated vimentin. LPS and heat-inactivated *Pseudomonas aeruginosa* (3×10^6^/mL) were positive controls. For the assay with anti-vimentin antibodies, neutrophils were simultaneously stimulated with CitVim at 1 µg/mL and various antibodies, namely; rabbit anti-vi (#ab45939, Abcam), Pritumumab (#MA5-41862, Invitrogen), mouse anti-cell surface vimentin (#H00007431-M08, Invitrogen) and anti-CitVim (#22054, Cayman Chemical) at a dilution range of 1:1,000 – 1:100 for 4 h. Mouse (Ms IgG) and rabbit (Rb IgG) isotype IgG control was used at 0.1 mg/mL. Cytokine and chemokine levels were assessed in the supernatant collected and centrifuged to remove remaining neutrophils.

### ROS formation and TLR4 inhibition

For the generation of reactive oxygen species (ROS), neutrophils were probed with DCFH-DA (2′-7′-dichlorofluorescein diacetate; 20 µM, Sigma-Aldrich) in culture medium untreated, or treated with vimentin, citrullinated vimentin and PMA for 4 h in the dark. The anti-TLR4 antibody (#ab13556, Abcam) at 1:100 dilution and TLR4 inhibitor (TAK-242, # 614316, Sigma-Aldrich) at 1 µM was added to the treated and untreated cells. The background fluorescence was determined for each condition and subtracted from the total fluorescence values before data analysis. The results were compared to the values in untreated condition (0), set as 1.

### Colocalization studies

We conducted confocal microscopy analysis to determine the surface localization of eVim, CitVim, and TLR4 in human neutrophils. Initially, 2×10^5^ neutrophils were cultured on poly-l-lysine-treated glass coverslips at 37°C. Subsequently, eVim and CitVim (at a concentration of 1 µg/mL) were introduced or omitted (CT) to the cells for a 1 h incubation period. After incubation, the cells were carefully washed with PBS, and the surface presence of eVim and CitVim was vitally stained using a mouse anti-Vim antibody (for eVim, #ab8069, Abcam) and mouse anti-CitVim antibody (for CitVim), both diluted at 1:500, for 1 hour at 4°C. Following vital staining, the cells were fixed with 3.7% paraformaldehyde (30 min, RT), blocked with 0.1% BSA (30 min, RT), and permeabilized with 0.1% Triton X100 (15 min, RT). Next, neutrophils were incubated with an anti-TLR4 antibody (1:500 dilution) for 1 h at RT. After washing off the primary antibody, secondary antibodies (1:500 dilution) were introduced to the cells for 1 hour at room temperature in the dark. Specifically, an anti-mouse antibody conjugated with AlexaFluor 488 was used for eVim and CitVim, while an anti-rabbit antibody conjugated with AlexaFluor 647 was employed for TLR4. Following a final wash, stained cells were mounted with an anti-fade solution and sealed with a glass coverslip. Subsequently, samples were scanned using a Stellaris 5 confocal microscope (Leica) with a 60× oil immersion objective. The degree of colocalization was quantitatively assessed using Mander’s coefficients through the JACoP plugin in FIJI ImageJ.

### Functionalization of AFM tips and substrates

Silicon nitride AFM tips (Bruker, MSCT-C) with a nominal spring constant of 0.01 N/m were employed to investigate the interaction between eVim/CitVim functionalized on cantilevers and TLR4 (#ab233665, Abcam) immobilized on a mica surface. All silicon nitride tips were functionalized following a modification of established protocols(29, 62). In brief, tips were silanized with 2% v/v 3-aminopropyltriethoxysilane (APTES, Sigma-Aldrich) in acetone for 10 min at room temperature (~22°C). Subsequently, tips were rinsed with deionized water (DI water) and immersed in 0.5% v/v glutaraldehyde in DI water for 30 min at 25°C. After another round of DI water rinsing, tips were incubated in 50 µL of eVim or CitVim solution at 100 µg/mL for 30 min at room temperature. Unbound proteins were washed away with DI water. To block any remaining aldehyde groups on the probe surface, the tip was treated with 100 µg/mL BSA in PBS for 1 min at room temperature. The tips were rinsed and then submerged in PBS buffer (to be used within 6 h).

For protein immobilization, mica surfaces were freshly cleaved and incubated with 0.01% poly-L-lysine overnight. The mica was washed with DI water and then incubated with 10 µL of recombinant TLR4 at 100 µg/mL for 30 min at room temperature. Unbound protein was washed off. A series of control experiments were conducted to assess the binding’s specificity. Specifically, tip AFM and mica surfaces were functionalized with BSA, or the protein addition step was omitted.

### AFM force measurements

AFM was employed to assess single-molecule binding interactions between eVim/CitVim and TLR4 on a mica surface, utilizing a NanoWizard 4 BioScience JPK Instruments Bruker atomic force microscope operating in force spectroscopy mode. The investigation of eVim/CitVim–TLR4 interactions involved recording AFM force– distance (FD) curves. TLR4 immobilized on mica surfaces was probed in PBS at room temperature. Up to 40 force maps, each comprising 16 × 16 points and corresponding to a scan area of 5 μm × 5 μm, were acquired for each sample, covering multiple random spots on mica or confluent cell surfaces. A set point of 0.15 nN was consistently applied for all measurements, and a 5 μm/s AFM cantilever approach/retraction speed was employed to probe TLR4 immobilized on mica.

For the analysis of specific adhesion events, a worm-like chain (WLC) model, offering reaction forces and loading rate (LR) data (processed using JPK data processing and the JPK built-in analysis software), was employed. The retraction part of the obtained curves was utilized to classify and determine eVim/CitVim and TLR4 interactions. Curves exhibiting specific adhesion displayed a distinctive “shape,” amenable to fitting using the WLC model. JPK software facilitated the WLC model fitting in the segment of the force curve, indicating polymer stretching, and corresponding loading rate (LR) values were simultaneously determined. Forces represent averages derived from all collected force–distance curves where adhesion forces were observed.

### Dot blot binding assay

Dot blot binding assays were conducted to validate the binding capabilities of recombinant Vim and CitVim proteins to TLR4, employing a modified approach based on previous methodologies (63, 64). Various quantities (1.0, 2.0, and 4.0 μg) of eVim, CitVim, and BSA were transferred in 10 µL volumes onto NC membranes with a pore size of 0.2 μm (Bio-Rad). Following the transfer, the membranes were dried and then blocked with 5% skim milk in Tris-buffered saline–Tween 20 (0.1%) (TBS-T) for 2 h at room temperature. Subsequently, the membranes were washed with TBS-T (3 x 15 min) and incubated with whole cell lysate of human neutrophils, equivalent to 5×10^6^ cells per blot, in TBS-T with gentle shaking overnight at 4°C. After extensive washing with TBS-T, the blots were probed with the respective primary antibodies (anti-Vim, anti-CitVim, or anti-TLR4) at a 1:500 dilution in TBS-T overnight at 4°C. Following another round of washing, the blots were incubated for 2 h at room temperature with an IRDye 800CW-conjugated goat anti-rabbit or goat anti-mouse secondary antibody at a dilution of 1:5,000. Images were visualized using the Odyssey LiCor Imaging System (LiCor Biosciences), and spot intensities were quantified using Image Studio Acquisition Software.

### Quantification and statistical analysis

Statistical analysis and data visualization were performed using Origin 2021 software. Gene-gene coexpression analysis using the STRING database and analysis of enriched pathways according to the Kyoto Encyclopedia of Genes and Genomes (KEGG) were analyzed using STRING v11.5 web software. Statistical comparisons were made using unpaired Student’s t-test (between two samples) and one-way ANOVA followed by Tukey’s post hoc test (among multiple samples). Differences were considered statistically significant at p < 0.05. We confirmed the validity of the biological replicates and enhanced the reliability of the data by repeatedly performing independent experiments using at least three independent samples. Data are generally presented as mean ± SD of n = x experiments, with x indicating in figure legends the number of independent experiments performed unless otherwise stated.

## Data availability

All data that support the findings of this study are available from the corresponding authors upon reasonable request.

## Abbreviations

AFM: Atomic force microscope
APTES: 3-aminopropyltriethoxysilane
BSA: Bovine serum albumin
cDNA: Complementary DNA
CitH3: citrullinated histone H3
CitVim: Citrullinated vimentin
Col: Collagen
COVID-19: Coronavirus disease 2019
CSV: Cell-surface vimentin
CTGF: Connective tissue growth factor
CTGF: Control, untreated condition
DAMP: Damage-associated molecular pattern
DCFH-DA: 2′-7′-dichlorofluorescein diacetate
DMEM: Dulbecco’s Modified Eagle’s Medium
DNA: Deoxyribonucleic acid
ECM: Extracellular matrix
eVim: Extracellular vimentin
FBS: Fetal bovine serum
FD: Force distance
FITC: Fluorescein isothiocyanate
FLS: Fibroblast-like synoviocytes
GAPDH: Glyceraldehyde 3-phosphate dehydrogenase
GM-CSF: Granulocyte-macrophage colony-stimulating factor
HUVECs: Human umbilical vein endothelial cells
ICAM-1: Intercellular adhesion molecule 1
ICU: Intensive care unit
IL: Interleukin
IPF: Idiopathic pulmonary fibrosis
KEGG: Kyoto Encyclopedia of Genes and Genomes
LPS: Lipopolysaccharide
LR: Loading rate
MAPK3: Mitogen-activated protein kinase 3
mEF: Mouse embryonic fibroblasts
MOI: Multiplicity of infection
MPO: Myeloperoxidase
NC: Nitrocellulose
NE: Neutrophil elastase
NET: Neutrophil extracellular traps
NF-κB: Nuclear factor Kappa B
NOX2/gp91phox: NADPH oxidase subunit 2
oxLDL: Oxidized low-density protein
PAD: Peptidyl arginine deiminase
PBS: Phosphate-buffered saline
PDMS: Polydimethylsiloxane
PFA: Paraformaldehyde
PGMEA: Propylene glycol methyl ether acetate
PMA: Phorbol 12-myristate 13-acetate
PRRs: Pathogen recognition receptors
PTM: Post-translational modification Reverse transcription-quantitative polymerase chain
qRT-PCR: reaction
RA: Rheumatoid arthritis
RIPA: Radioimmunoprecipitation assay buffer
RNA: Ribonucleic acid
ROS: Reactive oxygen species
RT: Room temperature
SD: Standard deviation
SDS-PAGE: Sodium dodecyl sulfate-polyacrylamide
TAK-242: Selective TLR-4 inhibitor
TGF-β: Transforming growth factor-β
TLR4: Toll-like receptor 4
TNF-α: Tumor necrosis factor-α
VEGF: Vascular endothelial growth factor
Vim: Vimentin
WGA: Wheat Germ Agglutinin
WLC model: Worm-like chain model

## Author contributions

Ł.S., K. F., P.A.G., A.E.P., P.A.J., and R.B. designed research; Ł.S., K.S., A.W., P.D., A.L., and S.O. performed research. Ł.S. and M.Z. contributed new reagents/analytic tools; Ł.S., P.D. analyzed the data; Ł.S. prepared figures; Ł.S. and R.B. wrote the paper.

## Ethics declaration

### Ethics approval and consent to participate

Procedures with human blood samples from healthy donors were performed in accordance with the approval of the Medical University of Bialystok ethics committee No. APK.002.8.2023. All donors signed an informed consent form, in accordance with the relevant ethics committee approval.

