## Supplemental information for "Extracellular Vimentin is a Damage-Associated Molecular Pattern Protein Serving as an Agonist of TLR4 in Human Neutrophils"

**Acknowledgment:** This work was supported by a National Science Center under a Preludium 21 grant (UMO-2022/45/N//NZ6/01454) awarded to Ł.S. and by the Medical University of Białystok to R.B. (B.SUB.23.326).

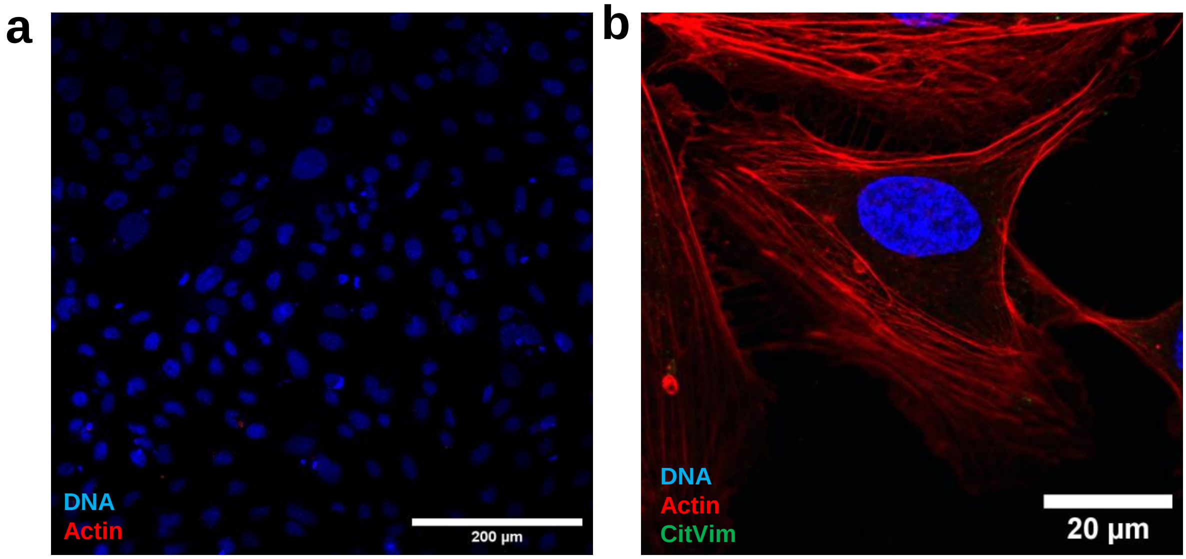


**Supplementary Figure 1. Vital staining does not permeabilize cells and thus does not allow the visualization of intracellular targets. Untreated endothelial cells are negative for Citrullinated Vimentin. a** Human umbilical vein endothelial cells are vitally stained with primary anti-β-Actin antibody (#A5441, Sigma-Aldrich) as a control for the permeability of the cells. After Vital staining with anti-β-Actin (1 h, 4°C) cells were fixed. Anti-mouse Alexa Fluor 647 was used as a secondary antibody.  Scale bar, ~200 µm. **b** HUVECs stained for citrullinated vimentin (green), actin (red), and nuclei (blue). Scale bar, ~20 µm.

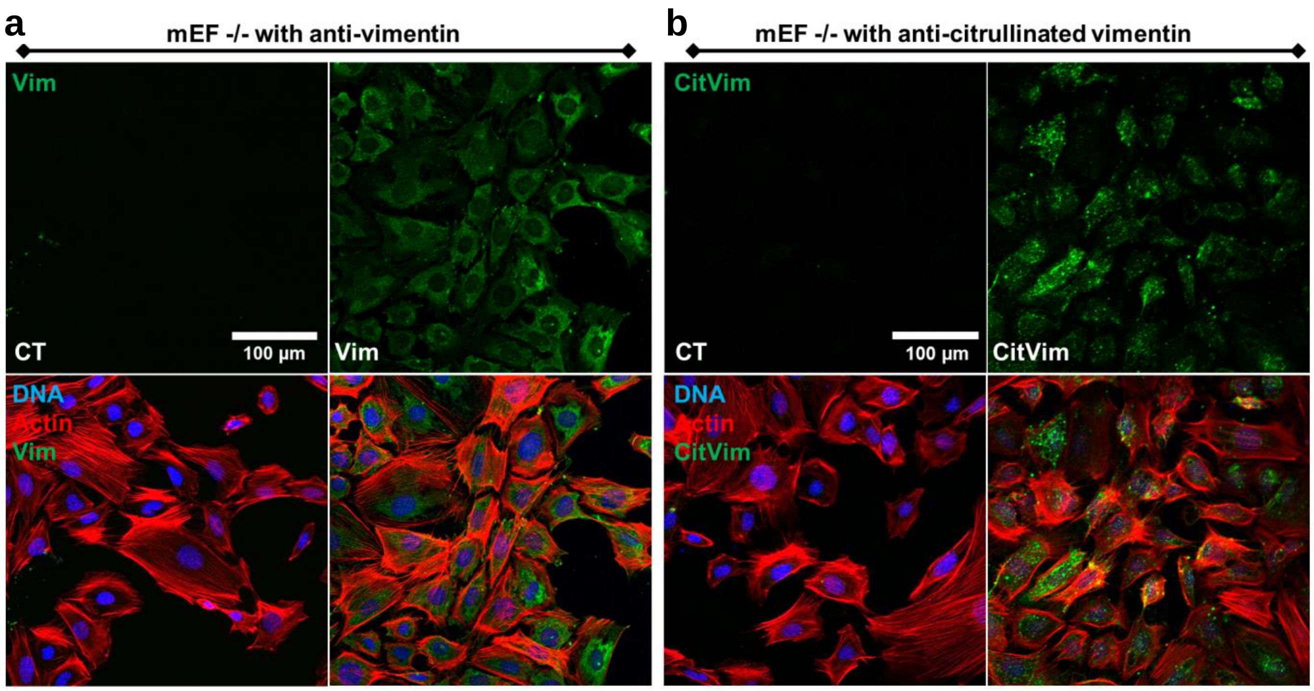


**Supplementary Figure 2.** Images of mouse embryonic fibroblasts lacking vimentin expression (mEF -/-) preincubated or not with vimentin (**a**) or citrullinated vimentin (**b**) for 1 h. Vimentin or citrullinated vimentin (green), actin (red), and nuclei (blue). Scale bar, ~100 μm.

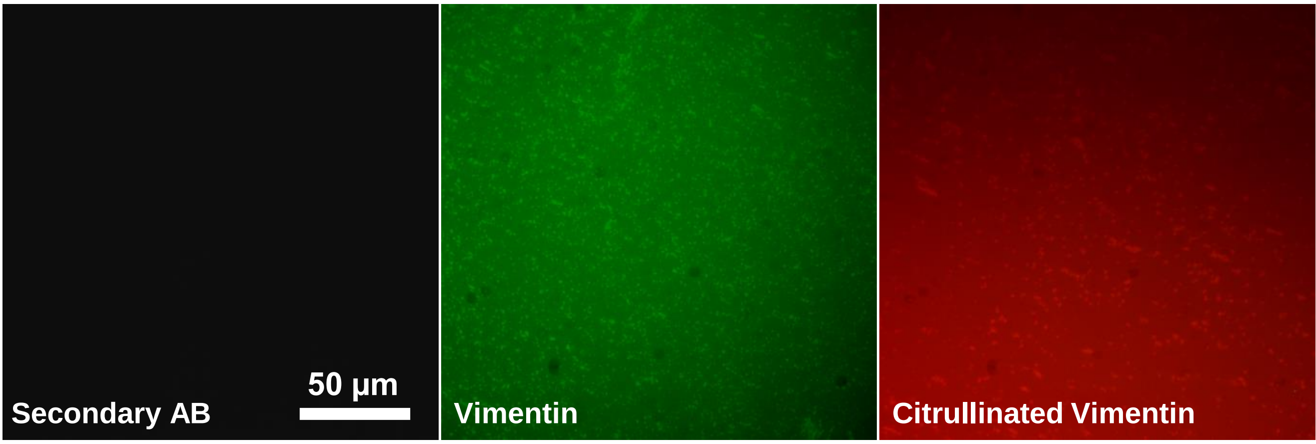


**Supplementary Figure 3. 30 kPa hydrogels uncoated (Secondary AB), coated with vimentin and citrullinated vimentin.** Vimentin was stained with rabbit polyclonal to Vimentin (Abcam, #ab45939). Citrullinated vimentin was stained with citrullinated vimentin mouse monoclonal antibody, Clone 12G11(Cayman Chemical, #22054). Secondary antibody – Goat anti-rabbit AlexaFluor 488 (green) and Goat anti-mouse AlexaFluor 647 (red). Scale bar, ~ 50 μm.

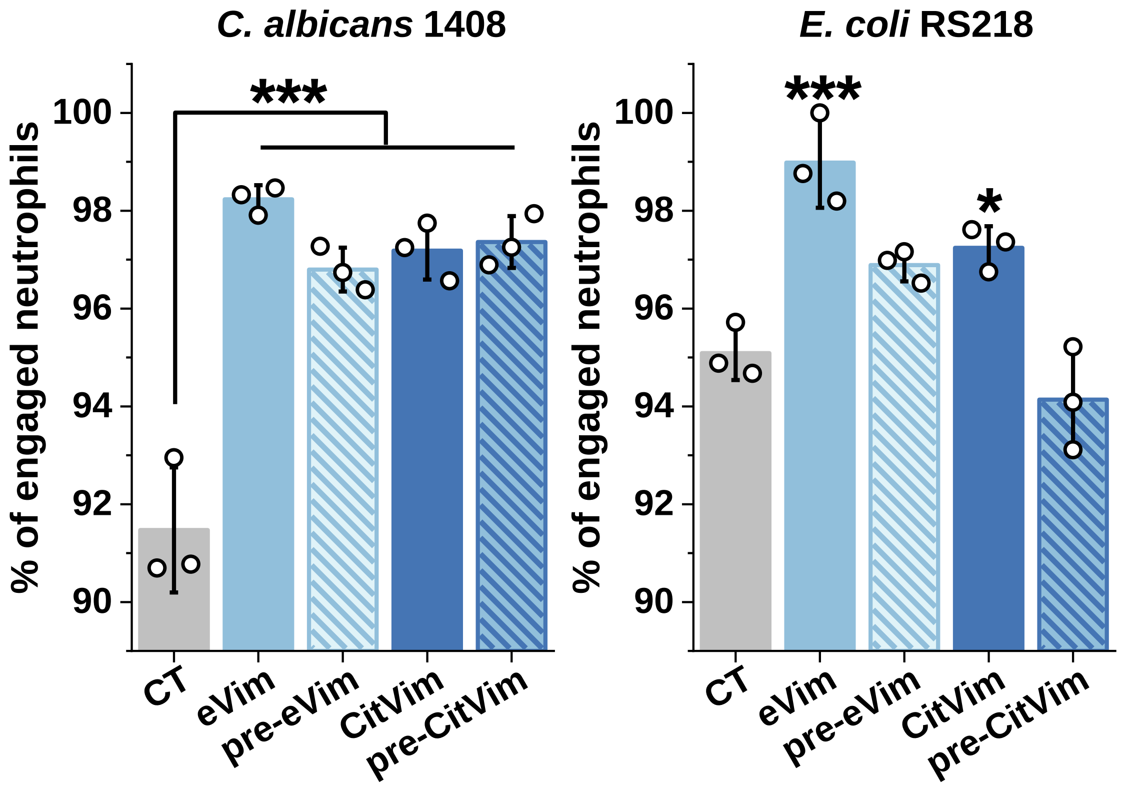


**Supplementary Figure 4.** Percentage uptake for vimentin (eVim – 1 μg/mL) or citrullinated vimentin (CitVim – 1 μg/mL) treated human neutrophils ingesting C. albicans cells (n = 3) at MOI of 10 and E. coli at MOI of 100. Additionally, neutrophils were preincubated with Vimentin (pre-eVim) and Citrullinated vimentin (pre-CitVim) at 1 μg/mL for 1 h in cell culture media, then yeasts/bacteria were added to the neutrophils for 2h. Neutrophils taking up at least one fungal cell were manually tracked to allow a quantitative analysis of percentage uptake during the 2 h coincubation period. Data are presented as the mean ± standard deviation of the mean. *, $P \leq 0.05$; **, $P < 0.01$; ***, $P < 0.001$. Significance was determined by one-way analysis of variance (ANOVA) with Tukey's test.

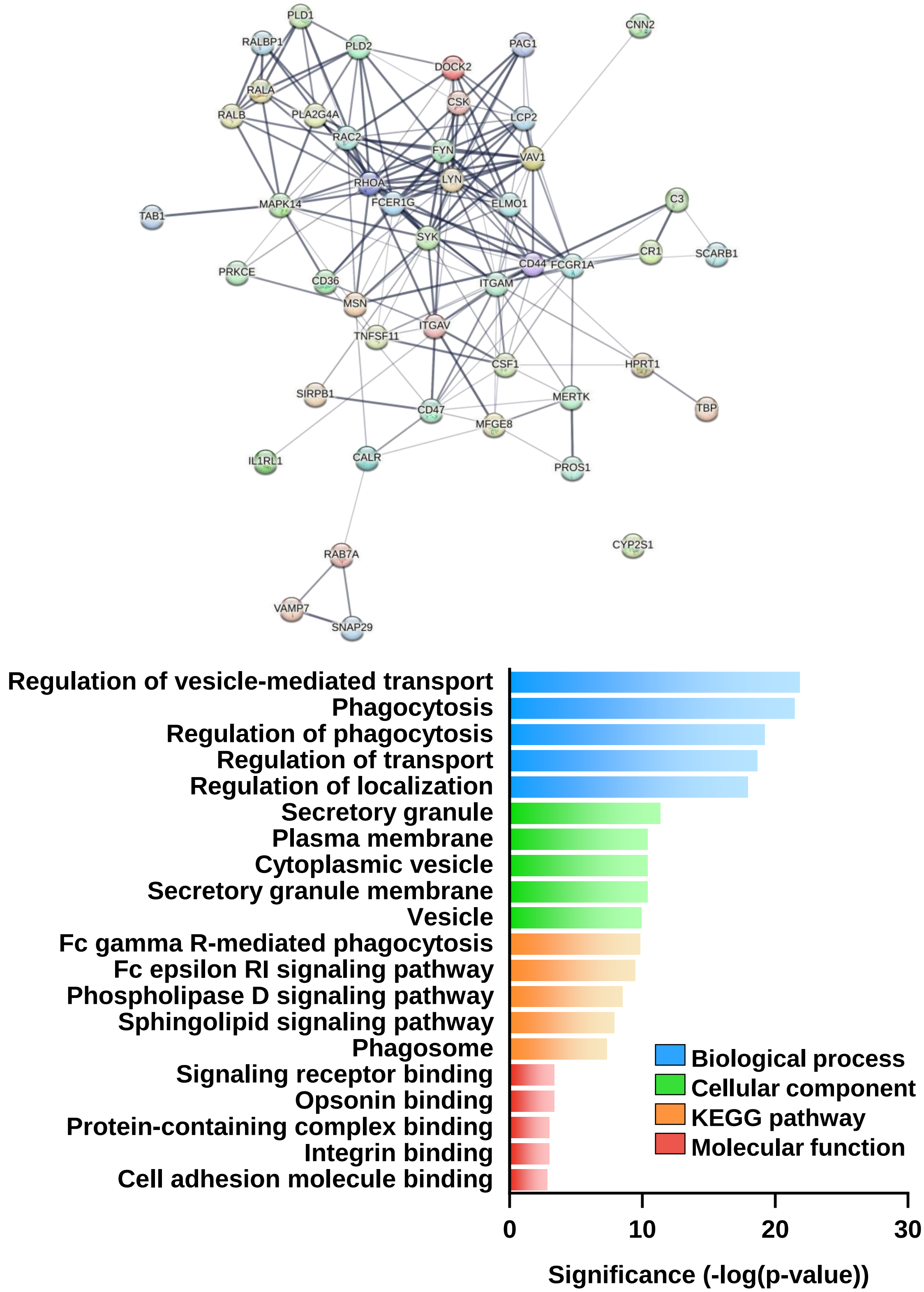


**Supplementary Figure 5.** The STRING network for human neutrophils stimulated with 1 μg/mL of vimentin for 2 h (upper panel). The edge thickness indicates functional and physical protein associations; the wider the edge, the stronger the association. Gene Ontology (GO) categories of the upregulated genes for human neutrophils stimulated with 1 μg/mL of vimentin for 2 h (bottom panel).

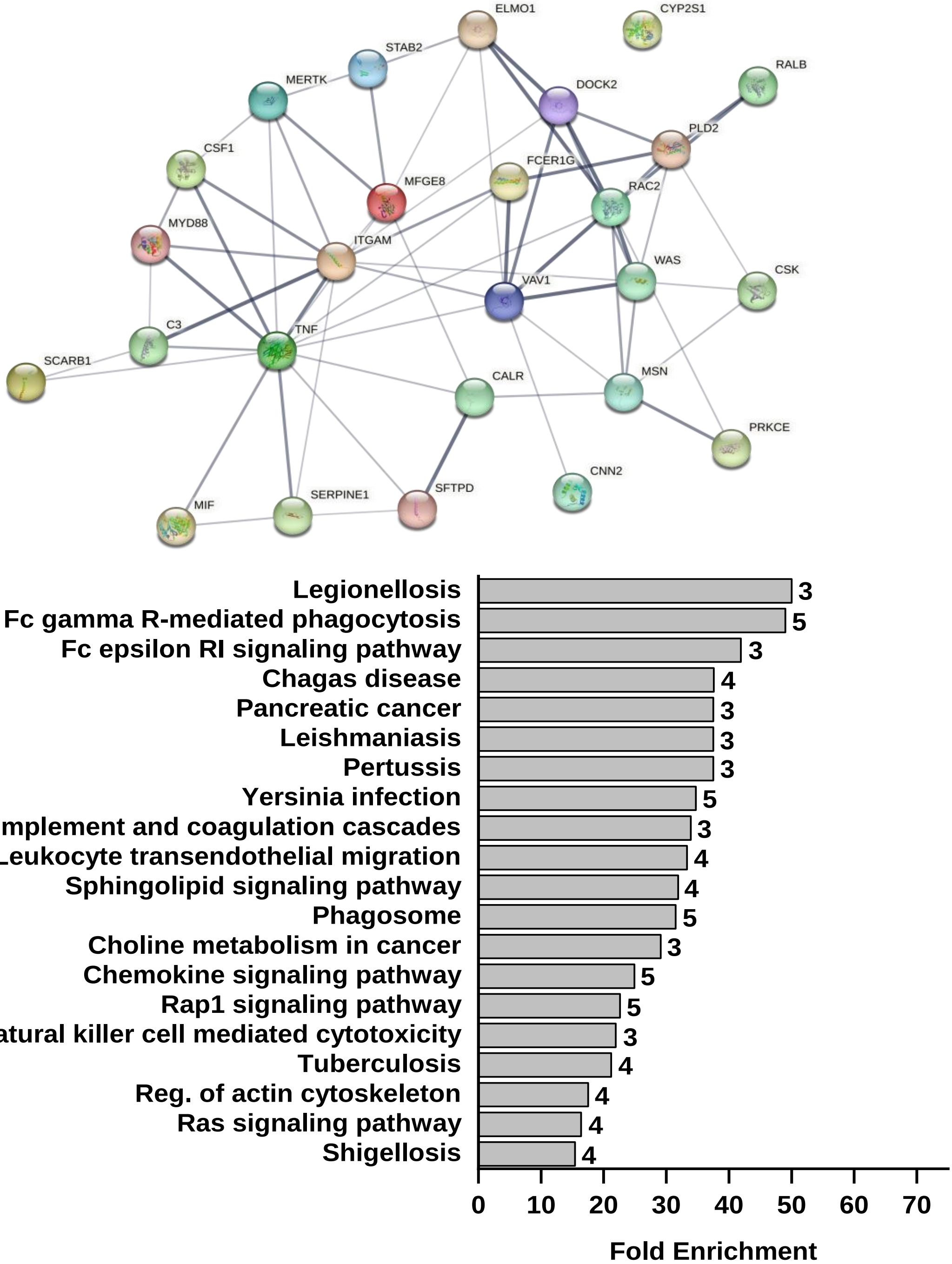


**Supplementary Figure 6. Phagocytosis-related genes association for neutrophils stimulated with citrullinated vimentin at 1 μg/mL.** The STRING network for human neutrophils stimulated with 1 μg/mL of citrullinated vimentin for 2 h (upper panel). The edge thickness indicates functional and physical protein associations; the wider the edge, the stronger the association. KEGG Pathways enrichment for human neutrophils stimulated with 1 μg/mL of citrullinated vimentin for 2 h (bottom panel). Too few genes were upregulated to determine their enrichment in biological processes, molecular function, or cellular components. Numbers at the end of each bar indicate the number of upregulated genes involved in a specific pathway.

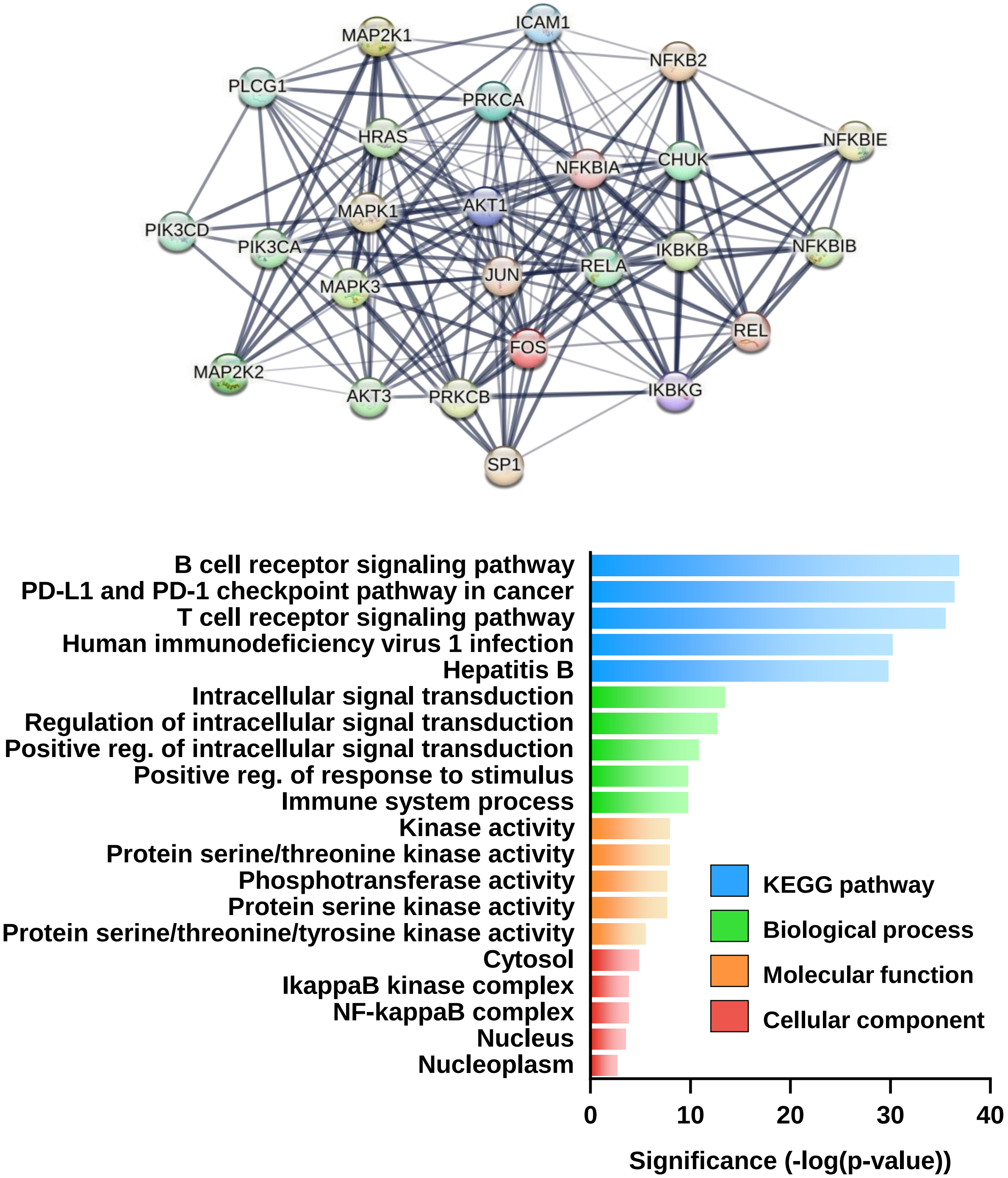


**Supplementary Figure 7.** The STRING network for human neutrophils stimulated with 1 μg/mL of citrullinated vimentin for 4 h (upper panel). The edge thickness indicates functional and physical protein associations; the wider the edge, the stronger the association. Gene Ontology (GO) categories of the upregulated genes for human neutrophils stimulated with 1 μg/mL of citrullinated vimentin for 4 h (bottom panel).

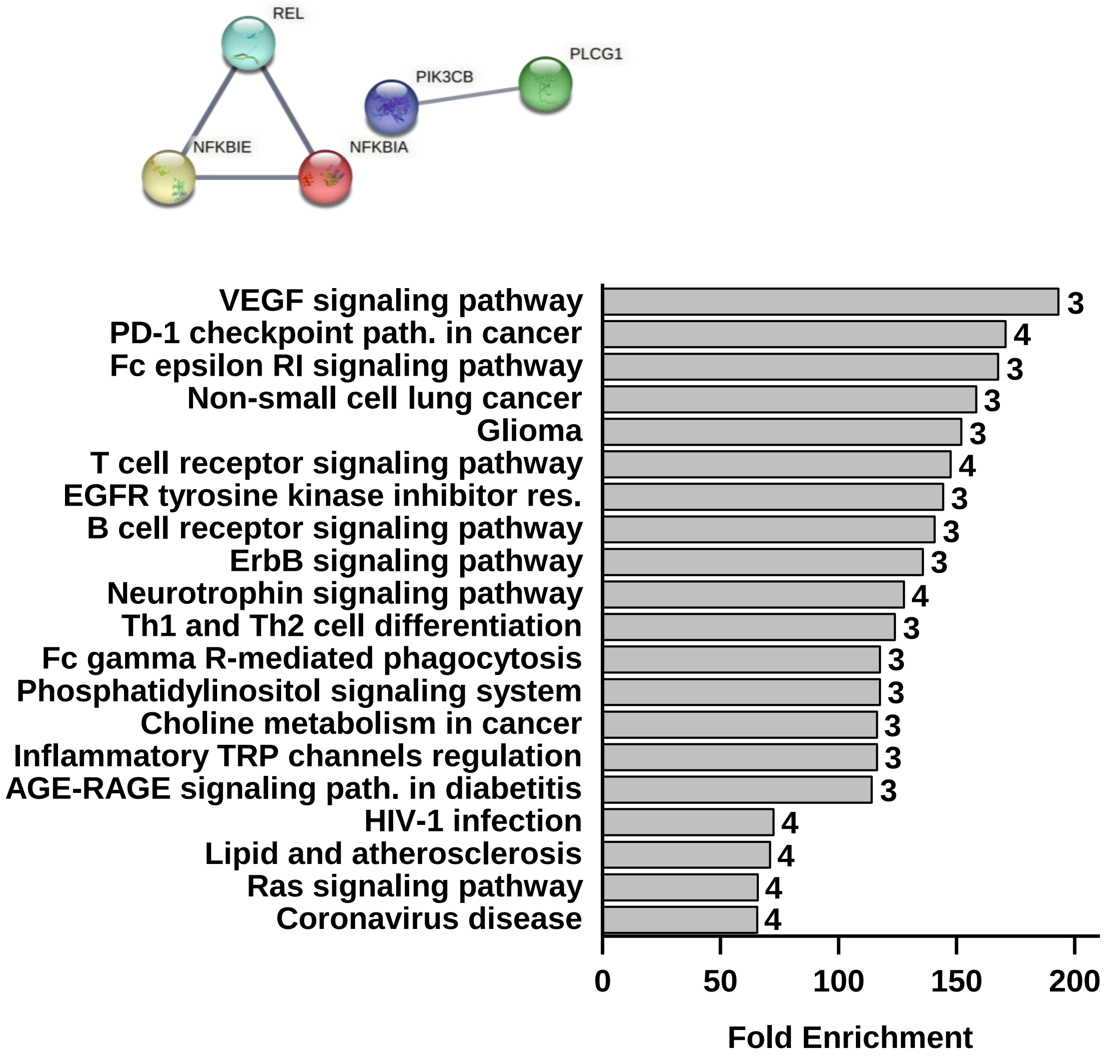


**Supplementary Figure 8. Inflammation-related genes association for neutrophils stimulated with vimentin at 1 µg/mL.** The STRING network for human neutrophils stimulated with 1 µg/mL of vimentin for 4 h (upper panel). The edge thickness indicates functional and physical protein associations; the wider the edge, the stronger the association. KEGG Pathways enrichment for human neutrophils stimulated with 1 µg/mL of vimentin for 4 h (bottom panel). Numbers at the end of each bar indicate the number of upregulated genes involved in a specific pathway. Too few genes were upregulated to determine their enrichment in biological processes, molecular function, or cellular components.

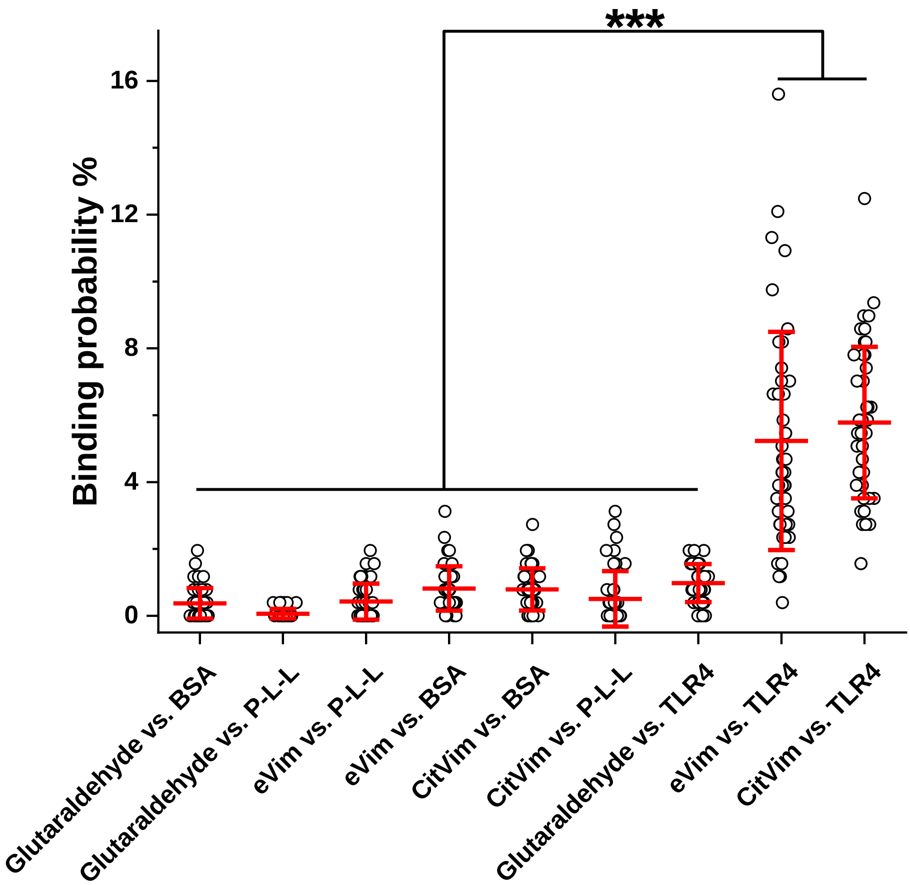


**Supplementary Figure 9. The binding of eVim and CitVim to TLR4 is specific in AFM settings.** Box plot of specific binding probabilities measured by AFM between tip functionalized with eVim, CitVim, or Glutaraldehyde only vs. mica with immobilized TLR4, bovine serum albumin (BSA), or Poly-L-Lysine (PLL). A single data point represents one force map and denotes the percentage of adhesive force curves in relation to all force curves acquired in this map ($n \geq 42$).

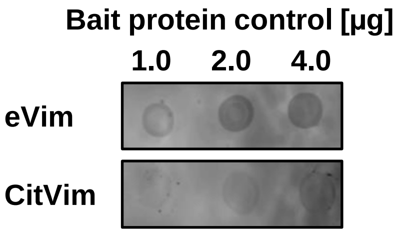


**Supplementary Figure 10.** To serve as a control in the dot blot binding assay, increasing amounts (1 to 4 µg) of eVim and CitVim were immobilized onto NC membranes and incubated with primary anti-Vim and anti-CitVim antibodies, subsequently detected using IRDye-conjugated secondary antibodies.

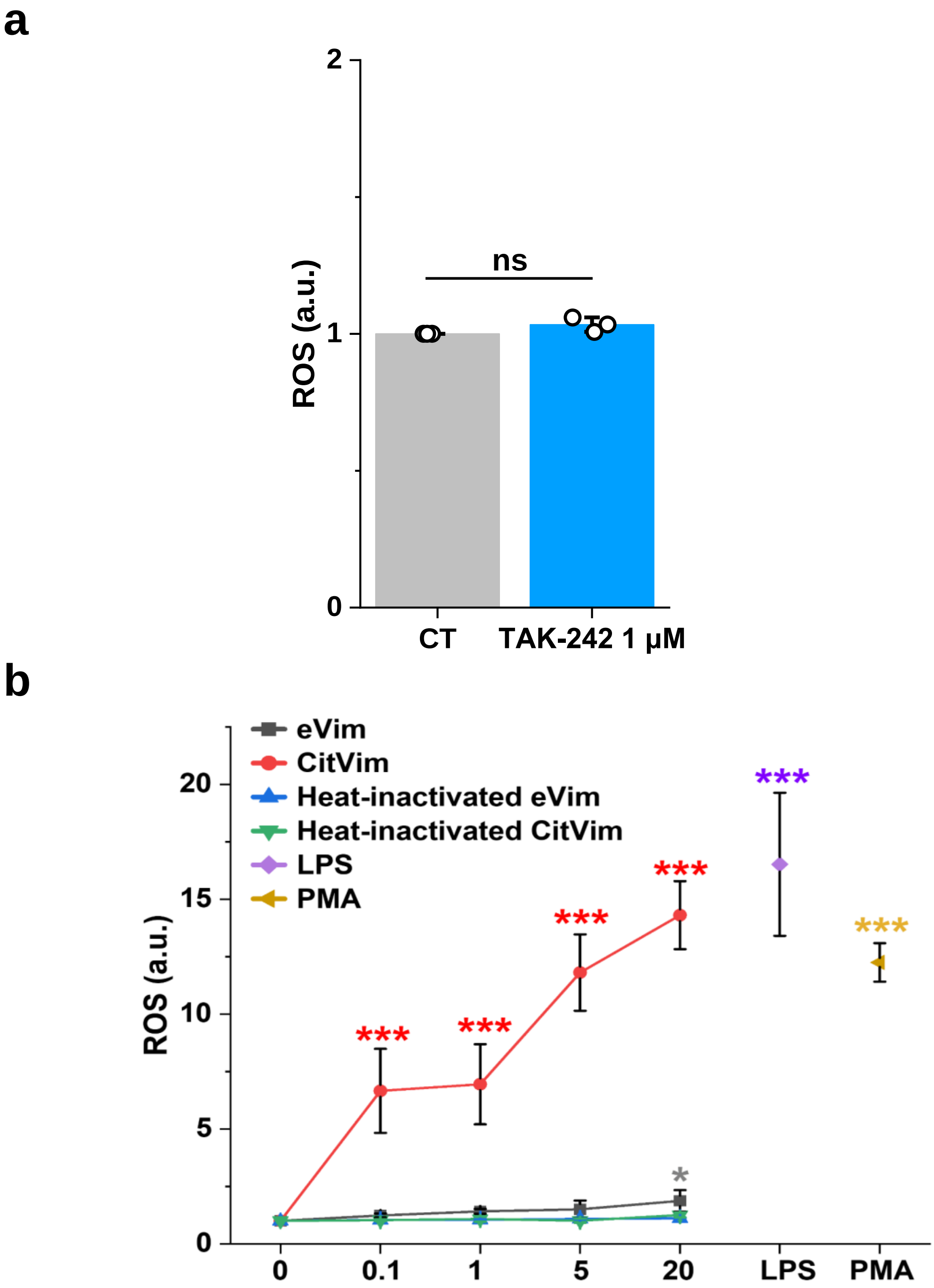


**Supplementary Figure 11. a. TLR4 inhibitor TAK242 at 1 µM does not influence ROS production compared to untreated neutrophils. b.** Release of reactive oxygen species (ROS) from human neutrophils was assessed after 4 hours of exposure to vimentin, citrullinated vimentin, heat-inactivated vimentin (0.1 – 20 µg/mL) and heat-inactivated citrullinated vimentin (0.1 – 20 µg/mL). LPS at 1 µg/mL and PMA at 100nM were used as a control. Data are presented as a normalized ROS production compared to the control condition (CT, 0) set as 1.0 with the mean ± SD (n = 3). Statistical analysis was performed using an unpaired Student's t-test (**a**) and ANOVA with post-hoc Tukey's test (**b**).

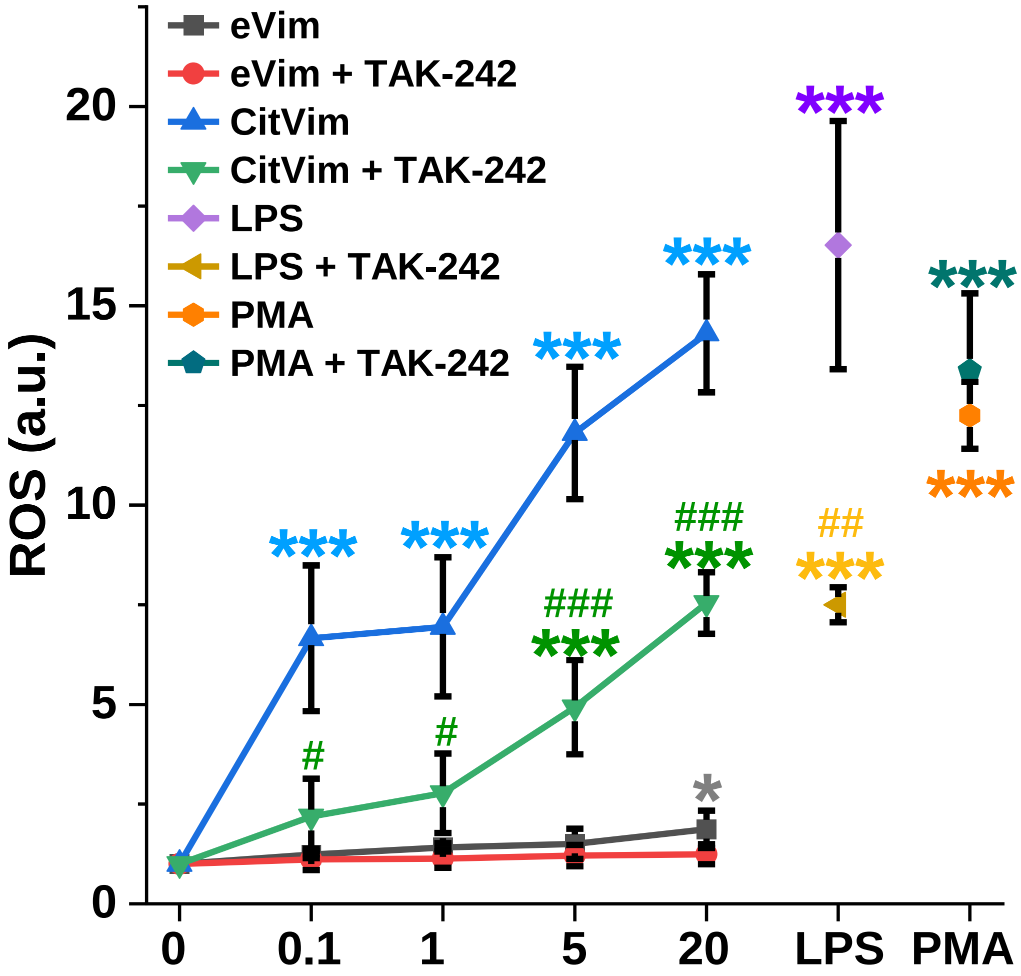


**Supplementary Figure 12.** Reactive oxygen species (ROS) release measured after exposing human neutrophils to varying concentrations of eVim, CitVim, LPS, and PMA, with and without TLR4 inhibition (TAK-242). ROS levels were normalized to untreated conditions (n = 3). Data are presented as the mean ± standard deviation of the mean. *, #, P ≤ 0.05; **, ##, P < 0.01; ***, ### P < 0.001. A hash sign (#) indicates a statistical significance comparison between corresponding eVim/CitVim and PMA compared to conditions with TLR4 inhibitor TAK-242 (i). Significance was determined by one-way ANOVA with Tukey's test or Student's t-test (f).

**Supplementary Table 1.** Antibodies and their epitopes used in Figure 7.

| Antibody | Cat. No. | Producer | Epitope |
|---|---|---|---|
| Rb anti-vim - polyclonal to the C-terminal part of vimentin | ab45939 | Abcam | Synthetic peptide containing 17 residues from within amino acids 425–466 of human vimentin. |
| Pritumumab – Recombinant monoclonal | MA5-41862 | Invitrogen | Cell-surface vimentin's C2 core region (coil 2 of the central rod). |
| Anti-CSV (Cell-surface vimentin) Clone 84-1 – polyclonal to vimentin | H00007431-M08 | Invitrogen | Human recombinant vimentin. |
| Anti-CitVim - Citrullinated Vimentin Monoclonal Antibody, Clone 12G11 | 22054 | Cayman Chemical | Synthetic peptide sequence from an internal region of human vimentin with citrullines at R144 and R146. |
